# Nanomaterials trigger functional responses in primary human immune cells

**DOI:** 10.1101/2024.10.17.618911

**Authors:** Vincent Mittelheisser, Olivier Lefebvre, Mainak Banerjee, Shayamita Ghosh, Amandine Dupas, Marie-Charlotte Diringer, Juliette Blumberger, Louis Bochler, Sébastien Harlepp, Annabel Larnicol, Angélique Pichot, Tristan Stemmelen, Anne Molitor, Chloé Moritz, Christine Carapito, Raphaël Carapito, Loïc J. Charbonnière, François Lux, Olivier Tillement, Jacky G. Goetz, Alexandre Detappe

## Abstract

Targeting the immune system with nanoparticles (NPs) to deliver immunomodulatory molecules emerged as a solution to address intra-tumoral immunosuppression and enhance therapeutic response. While the potential of nanoimmunotherapies in reactivating immune cells has been evaluated in several preclinical studies, the impact of drug-free nanomaterials on the immune system remains unknown. Here, we characterize the molecular and functional response of human NK cells and pan T cells to a selection of five NPs that are commonly used in biomedical applications. After a pre-screen to evaluate the toxicity of these nanomaterials on immune cells, we selected ultrasmall silica-based gadolinium (Si-Gd) NPs and poly(lactic-*co*-glycolic acid) (PLGA) NPs for further investigation. Bulk RNA-sequencing and flow cytometry analysis showcase that PLGA NPs trigger a transcriptional priming towards activation in NK and pan T cells. While PLGA NPs improved NK cells anti-tumoral functions in cytokines-deprived environment, Si-Gd NPs significantly impaired T cells activation as well as functional responses to a polyclonal antigenic stimulation. Altogether, we identified PLGAs NPs as suitable and promising candidates for further targeting approaches aiming to reactivate the immune system of cancer patients.

## Introduction

Over the past decade, immunotherapies became increasingly attractive for cancer treatment^[1–4]^. Among them, adoptive cell therapy (ACT) emerge as one of the most promising approaches^[5]^. These therapies aim to modify *ex vivo* the immune cells to induce the expression of an exogenous antigen receptor, enabling the detection of tumor antigens and hence resulting in potent anti-tumoral responses. T cells isolated from blood or tumor tissue were the first genetically reprogrammed immune cells for ACT^[6–8]^. The infusion of autologous modified T cells demonstrate positive clinical outcomes in relapsed hematological malignancies^[9,10]^, yet, adoptively transferred T cells are often associated with severe life-threatening side effects such as graft-versus-host disease, cytokine release syndrome (CRS), and neurological toxicities^[11]^. To overcome these limitations, emerging strategies utilizing the cytotoxic activity of non-MHC restricted NK cells are being evaluated clinically^[12]^. Allogeneic engineered NK cells administered to patients have shown tremendous potential to treat cancer patients, as they do not induce the abovementioned limitations while offering the possibility to be used off-the shelf^[13,14]^. In contrast to hematological malignancies, intratumoral infiltration of immune cells is restricted in solid tumors that quickly build an immunosuppressive TME which induces a rapid functional impairment of immune cells^[15–19]^. The emergence of cancer immunotherapies approaches allows to tackle the tumor microenvironment (TME) immunosuppression, but these therapies are associated with low response rate and life-threatening immune-related adverse events ^[20–23]^. Consequently, improving cancer immunotherapy efficacy and safety remains a priority.

One of the most promising solution lies in the utilization of nanoimmunotherapy approaches to specifically target immune cells and to promote immune cells mediated anti-tumoral responses through controlled spatiotemporal immunomodulation^[24]^. Specifically, nanoimmunotherapies were evaluated by using carbon nanotubes (CNTs) or a combination of CNTs and poly(lactic-*co*-glycolic acid) (PLGA) polymeric nanoparticles (NPs) to act as artificial antigen presenting cells, which are pivotal in ACT manufacturing^[25,26]^. In addition, polymeric NPs are also leveraged to deliver immune reactivating agents *in vivo*^[27]^ and to genetically engineer immune cells through *ex vivo* and *in vivo* virus-free transfection methods^[28,29]^. More recently, targeted ultrasmall metallic NPs were developed to monitor *in vivo* cancer expression of immune checkpoint molecules^[30]^. While these nanoimmunotherapies are successfully evaluated in preclinical trials, the impact of the NPs chemistry on primary immune cells remains surprisingly unexplored despite their known passive internalization.

Here, we use untargeted and drug-free NPs to precisely investigate the impact of their material on innate NK cells and adaptive pan T cells sourced from healthy donors to explore the fate of NPs post-internalization in immune cells, to ensure their safety profile and guide the selection of specific NPs based on the therapeutic need. To conduct pre-screening studies integrating cytotoxicity assessments and comprehensive proteomic analysis, we selected five nanosystems from the broad array being explored in both pre-clinical and clinical research. First, we included theranostic ultrasmall polysiloxane-based gadolinium nanoparticles (Si-Gd), which are currently evaluated in two Phase II clinical trials as magnetic resonance (MR)-guided radio-enhancers (NCT04789486, NCT04899908). To assess the potential impact of chelated metals on their behavior, we also evaluated a variant of these nanoparticles where gadolinium is substituted with terbium (Si-Tb). In addition, we explored inorganic terbium NPs (Tb NPs), leveraging their luminescent properties for fluorescence imaging and biological labeling, while also investigating whether metal chelation differentially affects their performance compared to unprotected metallic NPs. Poly(lactic-*co*-glycolic acid) (PLGA) NPs were included for their well-established role as multifunctional drug carriers. Finally, carbon nanotubes (CNTs) were selected as model non-organic drug delivery systems, given their distinctive structural and physicochemical characteristics. Based on these results, we selected two NPs with minimal cytotoxicity and effects on the proteome. We further evaluated their internalization in primary immune cells and highlight efficient internalization of the two NPs in both primary NK and pan T cells, with CD4^+^ T cells displaying higher internalization potential when compared to CD8^+^ T cells. Proteogenomic mapping of immune cells response to the selected nanomaterial revealed that PLGA NPs trigger pro-inflammatory transcriptional programs in NK and pan T cells, while ultrasmall polysiloxane-based gadolinium NPs (Si-Gd NPs) do not alter their transcriptomes. We further probed the effect of NPs on the functional behavior of immune cells and observed that PLGA NPs transcriptional priming of NK cells improved anti-tumoral function of NK cells. Conversely, Si-Gd NPs treatment impaired pan T cells responses to polyclonal antigenic stimulation. This study provides a unique resource of the impact of NP materials on human NK and pan T cells that will be beneficial to the tailoring of nanoimmunotherapeutic strategies in oncology.

## Results and Discussion

### Unequal internalization and subsequent toxicities of nanomaterials on human innate and adaptative lymphocytes

We use a library of five NPs, composed of (i) oxidized carbon nanotubes (oxCNTs), (ii) ultrasmall polysiloxane-based gadolinium (Si-Gd) NPs synthesized by a top-down method from core (gadolinium oxide) shell (polysiloxane) NPs^[31]^ and (iii) ultrasmall polysiloxane-based terbium (Si-Tb) NPs synthesized with a bottom-up one pot synthesis^[32]^, as well as (iv) unfunctionalized terbium fluoride (Tb) NPs and (v) PLGA NPs (**Fig.1A**). These NPs are used at both pre-clinical and clinical stages^[33]^ and cover a large range of hydrodynamic diameter distributions (from 5 to 120 nm) and ζ-potentials (from −50 to 40 mV), representing a wide spectrum of the nanomedicine field (**Fig.S1A−N**). To assess the impact of these NPs on primary human immune cells, we negatively isolate NK and pan T cells from healthy donor buffy coats with high purity (>95%) and subject them to a cytotoxicity-driven screening test (**Fig.S2A−B**). We incubate the sorted NK and pan T cells with increasing NPs concentration for 48 h (**Fig.1B**). We observe that oxCNTs markedly decrease NK cells (IC_50_ = 68.37µg/mL) and pan T cells (IC_50_ = 47.65µg/mL) viability at low concentrations. Inversely, the other tested nanomaterials do not significantly decrease immune cells viability (**Fig.1B**). We perform a whole proteome (liquid chromatography-tandem mass spectrometry, LC-MS/MS) analysis to further assess nanomaterials impact on immune cells (**Fig.S3A**). While terbium-based NPs (Si-Tb and Tb NPs) do not overtly affect immune cell viability (**Fig.1B**), the whole proteome analysis reveals an upregulation of proteins linked with necroptosis (**Fig.S3A**). Based on this initial pre-screening that combines proteomic and viability read-outs, we exclude cytotoxic oxCNT and terbium-based NPs as they appear unattractive for further preclinical development in this context. On the other hand, Si-Gd and PLGA NPs display no toxicity towards primary human immune cells (IC_50_ > 1 mg/mL) (**Fig.1B**) and the observed deregulated proteins are not associated with specific gene ontology terms (**Fig.S3A**). We thus decided to further evaluate their impact on immune cells using concentrations of 150 µg/mL of Si-Gd NPs^[34]^ and 50 µg/mL of PLGA NPs^[35]^ (median of the tested range for immune cells viability assay). We next confirm that Cyanine5.5-labelled Si-Gd NPs and PLGA NPs efficiently penetrate NK and pan T cells despite their low phagocytic potential (**Fig.1C**). Furthermore, we assess their cellular uptake via flow cytometry 48h post-treatment. We observe that Si-Gd NPs and PLGA NPs are uniformly internalized in NK cells, forming a single continuous population with no significant differences between CD56^High^ and CD56^Low^ NK cells (**Fig.1D-E**). Conversely, we notice that pan T cells exhibited two distinct populations based on the uptake levels - *i.e.* one with low internalization rates comparable to NK cells and another with significantly higher levels of both NPs (**Fig.1D**). Further flow cytometric analysis revealed that high internalizing T cells population predominantly comprises CD4^+^ T cells, while the low internalizing population is enriched in CD8^+^ T cells (**Fig.1F**). Importantly, increasing the concentration of NPs resulted in a dose-dependent uptake in NK cells and low-internalizing pan T cells (**Fig.S3B**). At the opposite, the high internalizing pan T cells population reached a plateau starting at the lowest tested Si-Gd NPs concentration and only a marginal increase in PLGA NPs uptake at higher doses (**Fig.S3C**).

**Figure 1.**
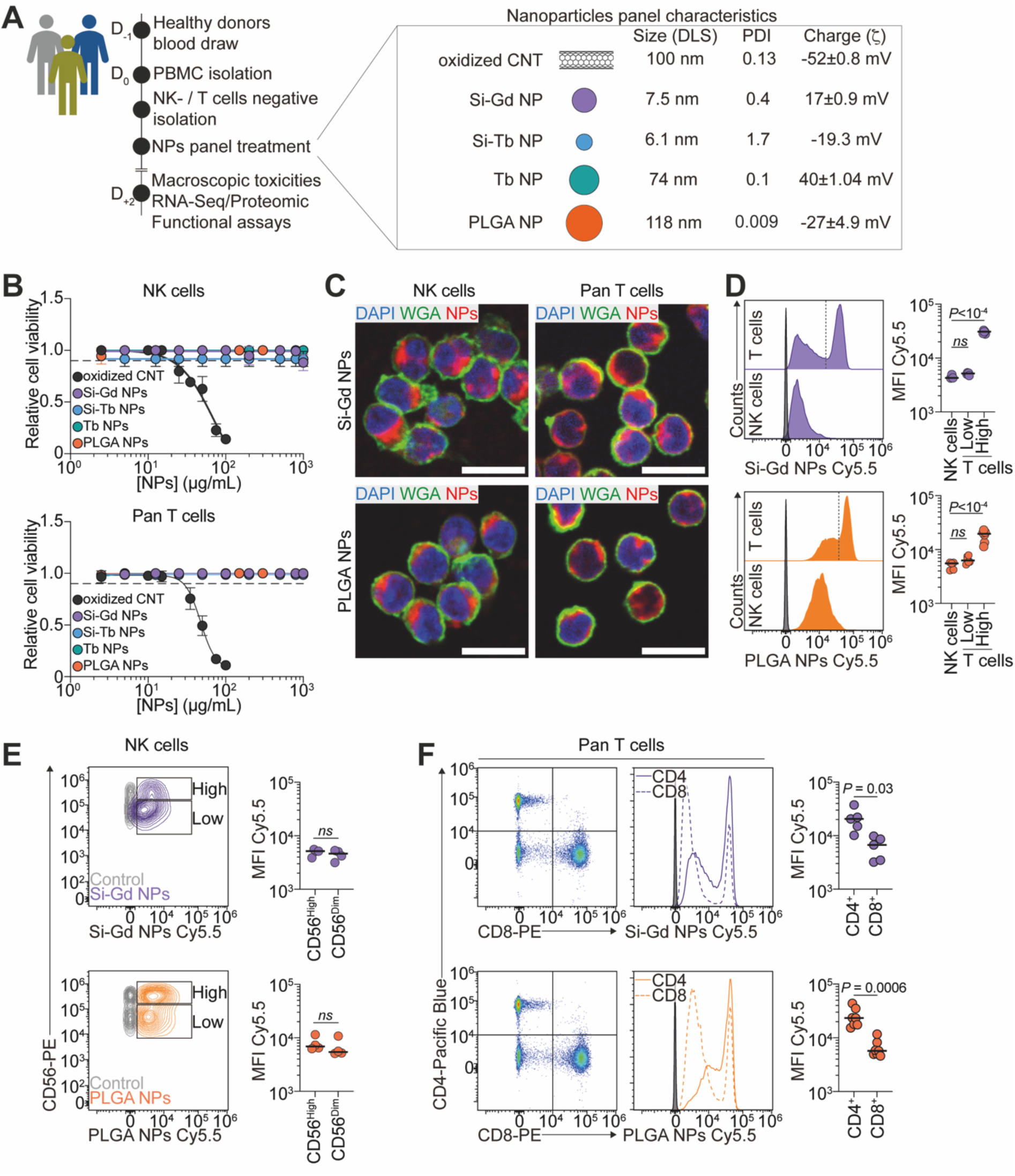
Nanomaterials internalization vary across human NK cells and pan T cells. **A**. Infographics illustrating the pipeline of the assessment of nanomaterial impact on primary human NK- and pan T cells. Nanomaterial physicochemical characteristics are displayed on the right. **B**. Nanomaterials impact on NK cells (upper panel) and pan T cells (lower panel) viability measured using CellTiterGlo assay. Data are representative of 3 independent experiments. Data are presented as mean ± s.d. **C**. Representative confocal micrographs of NK cells and pan T cells showing internalized Si-Gd NPs and PLGA NPs after 4h co-incubation at 37°C. In green Wheat Germ Agglutinin (WGA), in red nanomaterial (Cyanine5.5-labelled), in blue nuclei (DAPI). Scale bar = 10µm. **D**. Flow-cytometry assessment of nanomaterials internalization in NK- and pan T cells after 48h co-incubation at 37°C. Left: Representative histograms of the relative internalization of Si-Gd NPs (upper panel) and PLGA NPs (lower panel). Grey histogram: untreated control. Right: Quantification of the mean fluorescent intensity signal of the Cyanine5.5-labelled nanomaterials. Data are representative of 3 to 6 independent experiments and analyzed by a One-way ANOVA test with original FDR method of Benjamini-Hochberg after assessment of their gaussian distribution by Shapiro-Wilk test. **E**. Flow-cytometry assessment of nanomaterials internalization in CD56^High^ and CD56^Low^ NK cells after 48h co-incubation at 37°C. Left: Representative flow cytometry contour plots of the relative internalization of Si-Gd NPs (upper panel) and PLGA NPs (lower panel). Right: Quantification of the mean fluorescent intensity signal of the Cyanine5.5-labelled nanomaterials. Data are representative of 4 independent experiments and analyzed by a Student’s t-test (Si-Gd NPs) or Mann-Whitney test (PLGA NPs) after assessment of their gaussian distribution by Shapiro-Wilk test. **F**. Flow-cytometry assessment of nanomaterials internalization in CD4^+^ and CD8^+^ pan T cells after 48h co-incubation at 37°C. Left: Representative flow cytometry contour plots of the relative internalization of Si-Gd NPs (upper panel) and PLGA NPs (lower panel). Grey histogram: untreated control. Right: Quantification of the mean fluorescent intensity signal of the Cyanine5.5-labelled nanomaterials. Data are representative of 5 to 7 independent experiments and analyzed by a Student’s t-test with Welch’s correction (Si-Gd NPs) or Mann-Whitney test (PLGA NPs) after assessment of their gaussian distribution by Shapiro-Wilk test.

Altogether, we demonstrate that Si-Gd NPs and PLGA NPs are successfully internalized by NK and pan T cells without associated toxicities. Yet, NK cells and CD4^+^ or CD8^+^ T cells are not equal with regards to nanomaterials uptake.

### PLGA NPs trigger a pro-inflammatory signature in immune cells

To provide a refined analysis of the impact of the NPs on human NK and pan T cells, we perform a gene expression analysis using bulk RNA sequencing after 48 hours of treatment. Unsupervised analysis of the entire transcriptome using principal component analysis (PCA) reveal that PLGA NPs-treated populations clustered separately from control (untreated) and Si-Gd NPs-treated cells in the principal component space, indicating significant transcriptomic differences (**Fig.2A**). We identify 249 differentially expressed genes (DEGs) in NK cells and 395 DEGs in pan T cells following PLGA NPs exposure (adjusted *p*-value ≤ 0.05 and fold-change > 2) (**Fig.S4A and Table S1-2**). In contrast and in agreement with the PCA analysis, only 16 and 11 genes are differentially expressed in Si-Gd NPs-treated NK cells and pan T cells respectively (**Fig.S4A and Table S1-2**).

**Figure 2.**
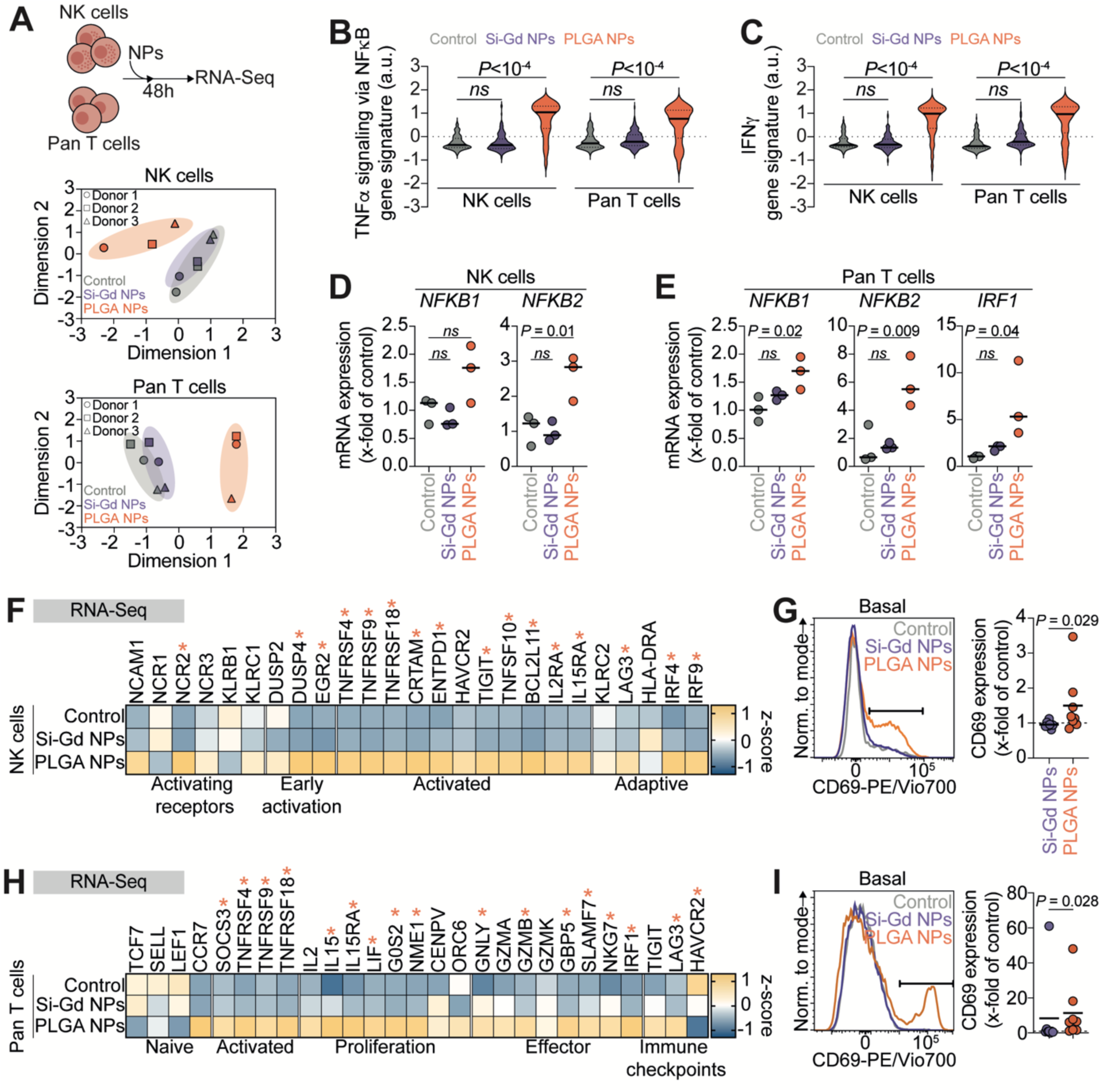
PLGA NPs transcriptionally prime immune cells activation. **A**. Transcriptomic impact of Si-Gd NPs and PLGA NPs treatment on immune cells. Upper panel: Schematic representation of the pipeline of the assessment of nanomaterial impact on primary human NK- and pan T cells transcriptome. Lower panel: Principal component analysis of the entire transcriptomic profile of NK cells and pan T cells untreated and treated with Si-Gd NPs or PLGA NPs. Each point represents one independent replicate. Axis labels represent the percent of variance as in the respective principal components (dimension 1 and dimension 2). **B-C**. Violin plots comparing aggregate expression distribution of genes related to TNFα signaling via the NFκB pathway (**B**) or related to IFNγ signaling (**C**) according to the MSigDB Hallmark 2024 database. The solid line within each violin represents the median and dotted lines represent quartiles. Data are analyzed by a Kruskal-Wallis test with original FDR method of Benjamini-Hochberg after assessment of their gaussian distribution by Shapiro-Wilk test. **D**. NK cells expression of *NKFB1* and *NFKB2* mRNA expression as fold change relative to untreated control calculated using the 2^−ΔΔCt^ method (housekeeping: *GAPDH*). Data are representative of 3 independent experiments and analyzed by One-way ANOVA test with original FDR method of Benjamini-Hochberg after assessment of their gaussian distribution by Shapiro-Wilk test. **E**. Pan T cells expression of *NKFB1*, *NFKB2* and *IRF1* mRNA expression as fold change relative to untreated control calculated using the 2^−ΔΔCt^ method (housekeeping: *GAPDH*). Data are representative of 3 independent experiments and analyzed by One-way ANOVA test with original FDR method of Benjamini-Hochberg after assessment of their gaussian distribution by Shapiro-Wilk test. **F**. Heat-map showing Z-score values for RNA expression of NK cells activation and function associated genes. Gene names are represented on the top while associations are represented on the bottom. Stars represent significant statistical difference between untreated control NK cells and PLGA NPs-treated NK cells. **G**. Flow-cytometry analysis of NK cells activation. Left: Representative histograms of CD69 expression at the NK cells surface in the different treatment conditions. Right: Quantification of CD69 expression as fold change relative to untreated control. Data are representative of 7 to 8 independent experiments and analyzed by a Mann-Whitney test after assessment of their gaussian distribution by Shapiro-Wilk test. **H**. Heat-map showing Z-score values for RNA expression of pan T cells activation and function associated genes. Gene names are represented on the top while associations are represented on the bottom. Stars represent significant statistical difference between untreated control pan T cells and PLGA NPs-treated pan T cells. **G**. Flow-cytometry analysis of pan T cells activation. Left: Representative histograms of CD69 expression at the pan T cells surface in the different treatment conditions. Right: Quantification of CD69 expression as fold change relative to untreated control. Data are representative of 8 independent experiments and analyzed by a Mann-Whitney test after assessment of their gaussian distribution by Shapiro-Wilk test.

Upstream regulators analysis of the deregulated genes following PLGA NPs treatment reveals that TNF was the most significant activated regulator (NK cells Z-score 7.79; pan T cells Z-score 5.354 and *p*-values of overlap were 2.82×10^-42^ and 4.60×10^-38^, respectively). Other regulators in the top five included IL1B, NFκB, and IFN, all of which were predicted to be activated upon PLGA NPs treatment (**Fig.S4B**). When we compared expected protein-protein interactions network among upregulated genes in PLGA NPs-treated pan T cells, we found a complex interaction between TNF, NFκB, and IFN in NK-(**Fig.S4C**) and pan T cells (**Fig.S4D**).

Using an aggregate expression analysis, we confirm the global upregulation of tumor necrosis factor-α (TNFα) signaling via the nuclear factor-κB (NFκB) and interferon gamma (IFNγ) pathways in PLGA NPs-treated NK cells compared with the untreated and Si-Gd NPs-treated populations (**Fig.2B-C**). Overexpression of key downstream regulators NFκB and IRF1 mRNA is further confirmed by reverse transcription quantitative polymerase chain reaction (RT–qPCR) (**Fig.2D-E**). In addition to significant enrichments of upregulated genes in GO terms/pathways associated with cytokines signaling and several immune-associated pathways, we observe that PLGA NPs treatment also induce an upregulation of immune cells specific transcriptional programs such as NK cells cytotoxicity against tumour cells, α/β T cells activation and T_H_1 immune response (**Fig.S4E**).

When probing transcriptional programs specific to immune cells activation, we observe that PLGA NPs treatment lead to a significant overexpression of genes linked to NK cells activating receptors (*NCR2*/NKp44), immediate-early response genes (*DUSP4*, and *EGR2*), and co-stimulatory receptors (*TNFRSF4*/OX-40, *TNFRSF9*/4-1BB, *TNFRSF18*/GITR, *CRTAM* and *TNFSF10*/TRAIL)^[36,37]^ (**Fig.2F and Table S3**). Correlatively, PLGA NPs-treatment induce a trend increase in the early activation marker CD69 expression at the basal level in NK cells, which we further confirm by flow cytometry (**Fig.2G**). PLGA NPs also induced an overexpression of the high-affinity interleukin-2 receptor (*IL2RA*/CD25) and NME1 (Fold Change: 2.46; *q*-value = 4.02×10^-4^) which are critical for NK cells activation and proliferation^[38]^ (**Table S3**). Additionally, PLGA NPs-treated NK cells displayed an upregulation of *IFNG* (Fold Change: 7.46; *q*-value = 4.67×10^-5^), *ICAM1* (Fold Change: 2.29; *q*-value = 2.86×10^-5^) transcripts required for NK cells killing of tumour cells^[13,39]^ as well as *XCL1* (Fold Charge: 2.15; *q*-value = 2.10×10^-6^) transcripts linked to cDC1 recruitment and orchestration of adaptive immunity by NK cells^[40]^ (**Table S3**). Transcriptional programs analysis also revealed that PLGA NPs treatment induced the upregulation of genes associated with a NK cells adaptative/memory phenotype (*LAG3*, *IRF4* and *BCL2L11*/BIM)^[41–43]^ (**Fig.2F and Table S3**). These NK cells also overexpressed *CCR7* (Fold change: 2.97; *q*-value = 8.63×10^-10^) and *CD83* (Fold change: 3.38; *q*-value = 1.07×10^-23^), genes associated with migratory and helper phenotypes leading to high ability to produce IFN-γ^[44]^.

Similarly, transcriptional programs analysis of PLGA NPs treated pan T cells present a positive z-score for genes related with activation (*TNFRSF4*/OX-40, *TNFRSF9*/4-1BB, *TNFRSF18*/GITR), proliferation (*IL15*/*IL15RA, LIF*), and cell division (*G0S2*, *NME1*)^[45,46]^ as well as effector functions (*GNLY*, *GZMB*, *SLAMF7*, *NKG7*)^[47,48]^ (**Fig.2H and Table S3**). Alongside, PLGA NPs-treated pan T cells displayed an upregulation of ATF-like transcription factors such as *BATF* (Fold Change: 2.38; *q*-value = 0.001) and *BATF3* (Fold Change: 2.34; *q*-value = 0.04), that are essential checkpoints for early effector T cells differentiation^[49,50]^. Furthermore, as highlighted in the differentially expressed term analysis (**Fig.S4C**), these effector-primed T cells displayed a T_H_1 transcriptional landscape (*STAT1*, *ADAM12*)^[51,52]^ as well as a reduction in T_H_2 transcription factors transcripts (*GATA3*) and cytokines (*IL12B*, *TNF*, *IL17F*, *XCL1, CCL3L1*, *CCL4L1*) known to prime and to be expressed during T_H_1/TH_17_-like pro-inflammatory response^[53–55]^ (**Table S3**). Similarly to NK cells and in accordance with the upregulation of activation/effector-associated genes, PLGA NPs treatment increased CD69 expression at the basal level in pan T cells as observed by flow cytometry (**Fig.2I**). We confirmed such priming with a nanomaterial-induced dose-dependent decrease in CD3 expression (**Fig.S5A-B**) and with an increase in pan T cells size (**Fig.S5C-D**), that are classically associated with their activation^[56]^. On the other hand, and as expected, Si-Gd NPs did not induce pan T cells basal activation (**Fig.S5A-E**). Besides, transcriptional activation of pan T cells by PLGA NPs was further underpinned by the induction of multiple gene families associated with canonical IFN response module (*IFIT1*, *IFIT2*, *IFIT3*, *IFITM1*, *IFITM2*, *IRF1*, *IFI44*, *IFI44L*, *STAT1*, *GBP1*, *GBP4*, *GBP5*)^[47,57,58]^ (**Table S2**). Of note, we also found a significant increase in the expression of immune checkpoint transcripts (*LAG3*, *HAVCR2*/TIM3) (**Fig.2I and Table S3**) that can be counteract by the upregulation of *EGR2* (**Table S3**) whose role was recently described in maintaining anti-tumor responses of exhausted T cells^[59,60]^.

Altogether, we identify a transcriptional priming in NK and pan T cells upon drug-free PLGA NPs treatment, but not with Si-Gd NPs through bulk RNA sequencing of immune cells. Flow cytometry analysis further exhibit that drug-free PLGA NPs exposure was sufficient to trigger NK and pan T cells activation.

### PLGA NPs enhance anti-tumor functions of NK cells

As PLGA NPs could transcriptionally prime NK cells, we sought to investigate whether they could functionally activate their tumor-lytic properties (**Fig.3A and Fig.S6A-B**). We co-culture NPs-treated NK cells with their prototypical targets, the K562 cells, over 4h with or without supplementing the pro-survival cytokine IL-15 to mimic tumor microenvironments that often lack pro-survival cytokines required by cytotoxic NK cells^[61]^ (**Fig.3A**). With IL-15, both Si-Gd NPs and PLGA NPs-treated NK cells successfully clear the K562 targets (**Fig.3B**). Without supplementary IL-15, only PLGA NPs-treated NK cells efficiently induce killing of 25% of the target cells (**Fig.3B**). Of note, NK cells anti-tumoral functions alterations observed in control NK cells or upon Si-Gd NPs treatment are not resulting from an impaired NK cells activation, as all conditions exhibited similar CD69 expression levels (**Fig.3C**). In addition, both NPs-treatment do not impair K562 target cells recognition, as observed through the measurement of available NKG2D receptor at the NK cells surface by flow cytometry (**Fig.3D**). The killing capacity of NK cells also requires efficient cytotoxic granules polarization at the immune synapse (IS). We next induce IS formation and assess the distance between mature perforin and the plasma membrane of NK-K562 synapses (**Fig.3E**), using stable expression of palmitoylated tdTomato (**Fig.S6C**). Without IL-15, PLGA NPs-treatment favor cytotoxic granules convergence at the IS, as measured by the reduction of granules distance to the IS (**Fig.3E**). Concomitantly, PLGA NPs-treatment increase the percentage of NK cells presenting a polarized IS (**Fig.3E**). In contrast, neither Si-Gd NPs treated, nor untreated NK cells present polarized IS (or reduced distance between perforin and the IS) (**Fig.3E**), suggesting that, as expected, PLGA NPs treatment induce efficient K562 cells lysis through an increase in NK cells perforin polarization. Coming back to our proteogenomic analysis, we highlight that PLGA NPs induce the upregulation of proteins associated with NK cells maturation (IKZF3)^[62]^, cell adhesion (VLA-1: ITGA1/ITGB1)^[63]^ and vesicle transport (STX11)^[64]^ (**Fig.S6D**). Conversely, Si-Gd NPs repress the expression of these proteins and stimulate the expression of proteins associated with NK cells inhibitory signaling (HLA-E and GSK3B)^[65–67]^ that are less or not expressed in PLGA NPs-treated NK cells (**Fig.S6D**). NK cell-induced cytotoxicity requires both polarization of the cytotoxic granules and their fusion with the plasma membrane, allowing the release of their cytotoxic content. We next monitor NK cells degranulation by dosing the cytotoxic cytokines TNFα^[68]^ and IFNγ^[69]^ in the supernatant after 4h of NK cells co-incubated with their target K562 cells. Increased cytotoxic granules polarization in PLGA NPs-treated NK cells is corroborated by a trend, yet non-significant, augmentation in TNF-α and IFN-γ concentrations when compared to control and Si-Gd NPs-treated NK cells (**Fig.3F**).

**Figure 3.**
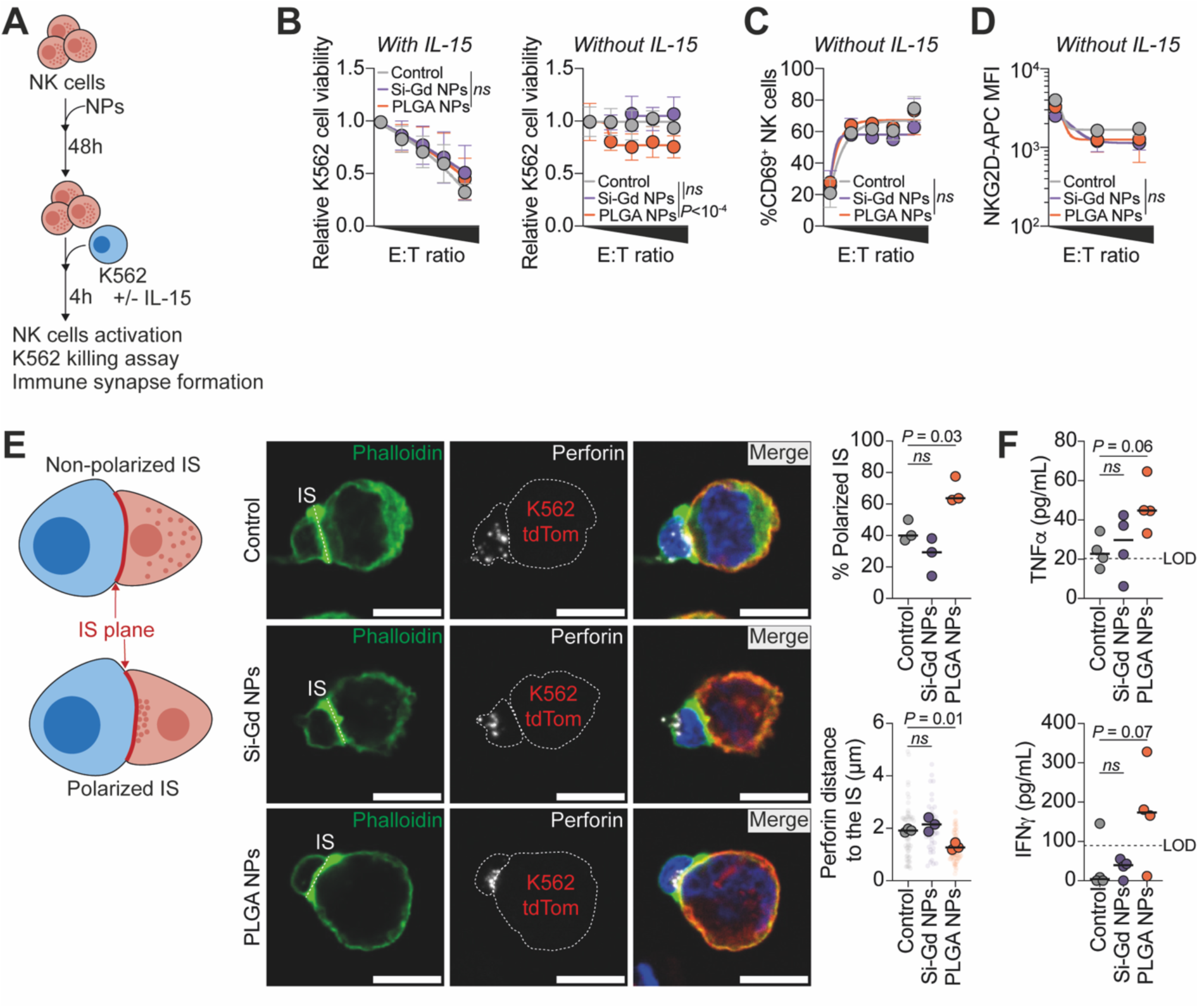
PLGA NPs enhance anti-tumor functions of NK cells by promoting perforin polarization at the immune synapse. **A**. Schematic representation of the pipeline of the assessment of Si-Gd NPs and PLGA NPs functional impact on primary human NK cells. **B**. Flow-cytometry assessment of K562 lysis induced by increasing nanomaterials-treated NK cells ratio (0.625:1; 1.25:1; 2.5:1 and 5:1) after 4h of co-incubation with 10ng/mL of IL-15 (left) or without (right). Data are representative of 6 to 8 independent experiments and analyzed by a Two-way ANOVA corrected with original FDR method of Benjamini-Hochberg. **C**. Percentage of CD69-positive NK cells assessed after co-incubation with K562 cells for 4h. Data are representative of 3 independent experiments and analyzed by a Two-way ANOVA corrected with original FDR method of Benjamini-Hochberg. **D**. Mean fluorescence intensity of NKG2D expressed at the NK cells surface after co-incubation with K562 cells for 4h. Data are representative of 3 independent experiments and analyzed by a two-way ANOVA corrected with original FDR method of Benjamini-Hochberg. **E**. Nanomaterials-treated NK cells immune synapse (IS) formation with K562. Left: Schematic representation of IS polarization and perforin distance evaluation. Center: Representative confocal micrographs of NK-K562 IS. In green Phalloidin-iFluor488, in white perforin dG9 (Alexa Fluor 647), in red K562 palmitoylated-tdTomato and in blue nuclei (DAPI). Scale bar = 10µm. Right: Percentage of polarized IS (upper panel) and mean perforin distance to the IS (lower panel). Data are representative of 3 independent experiments and analyzed by a One-way ANOVA test with original FDR method of Benjamini-Hochberg after assessment of their gaussian distribution by Shapiro-Wilk test. **F**. Concentration of TNFa (upper panel) and IFNg (lower panel) released in the supernatant after 4h of NK cells co-incubation with K562 cells at a 2:1 ratio. Data are representative of 4 independent experiments and analyzed by a One-way ANOVA test with original FDR method of Benjamini-Hochberg (TNFa) or by a Kruskal-Wallis test (IFNg) after assessment of their gaussian distribution by Shapiro-Wilk test. LOD: limit of detection.

According to these results, we demonstrate that NK cells transcriptional priming towards an activated phenotype induced by PLGA NPs results in an enhancement of their anti-tumoral cytotoxic capacity *in vitro* in a cytokines-deprived (*i.e.*, immunosuppressive) environment.

### Silica-based gadolinium NPs treatment impairs pan T cells functions

When investigating the impact of NPs on T cell functions, we notice from the unbiased proteomic approach previously described (**Fig.S3A**) that proteins involved in TCR signaling are significantly downregulated (CD48, PLCG1 and CARD11)^[70–72]^ upon Si-Gd NPs treatment (**Fig.4A**). These modifications are accompanied by a significant reduction in proteins associated with pan T cells cytoskeletal remodeling, including small GTPase (CDC42, RAC1 or SEPT7)^[73,74]^, actin binding proteins (Ezrin, EZRI or WAVE2)^[75,76]^ as well as formins (formin-like-1, FMNL1)^[77]^ (**Fig.4A**). Si-Gd NPs-treated pan T cells also display a significant decrease in the levels of integrin β1, which sustains cytotoxic T cells functions^[78,79]^ and their efficient intra-tumoral infiltration^[80]^. Similarly, we also observe a significant diminution in proteins required for terminal transport (KLC1 and KIF5B)^[81]^ and membrane fusion (SNAP23 and VAMP8)^[82,83]^ of lytic granule at the immune synapse (**Fig.4A**). Incidentally, PLGA NPs have no or little effects on these proteins (**Fig.4A**). Building on these observations, we aim at testing the effect of NPs on pan T cells function. We mimic stimulation of the TCR and co-receptors by coating surfaces with activating antibodies against CD3 (part of the TCR complex) and CD28 (a costimulatory receptor) and measure how T cells activate and spread, as they would over the surfaces of antigen presenting cells (APCs)^[84]^ (**Fig.4B**). We first observe that both Si-Gd and PLGA NPs treatment do not prevent global pan T cell activation when comparing CD69 expression after 24h of culture on a surface coated with increasing concentrations of anti-CD3 and a fixed concentration of anti-CD28 antibodies (hereafter, anti-CD3/CD28) (**Fig.4C**). This result further suggests that transcriptional priming is antigen-independent and mediated by intracellular cues.

**Figure 4.**
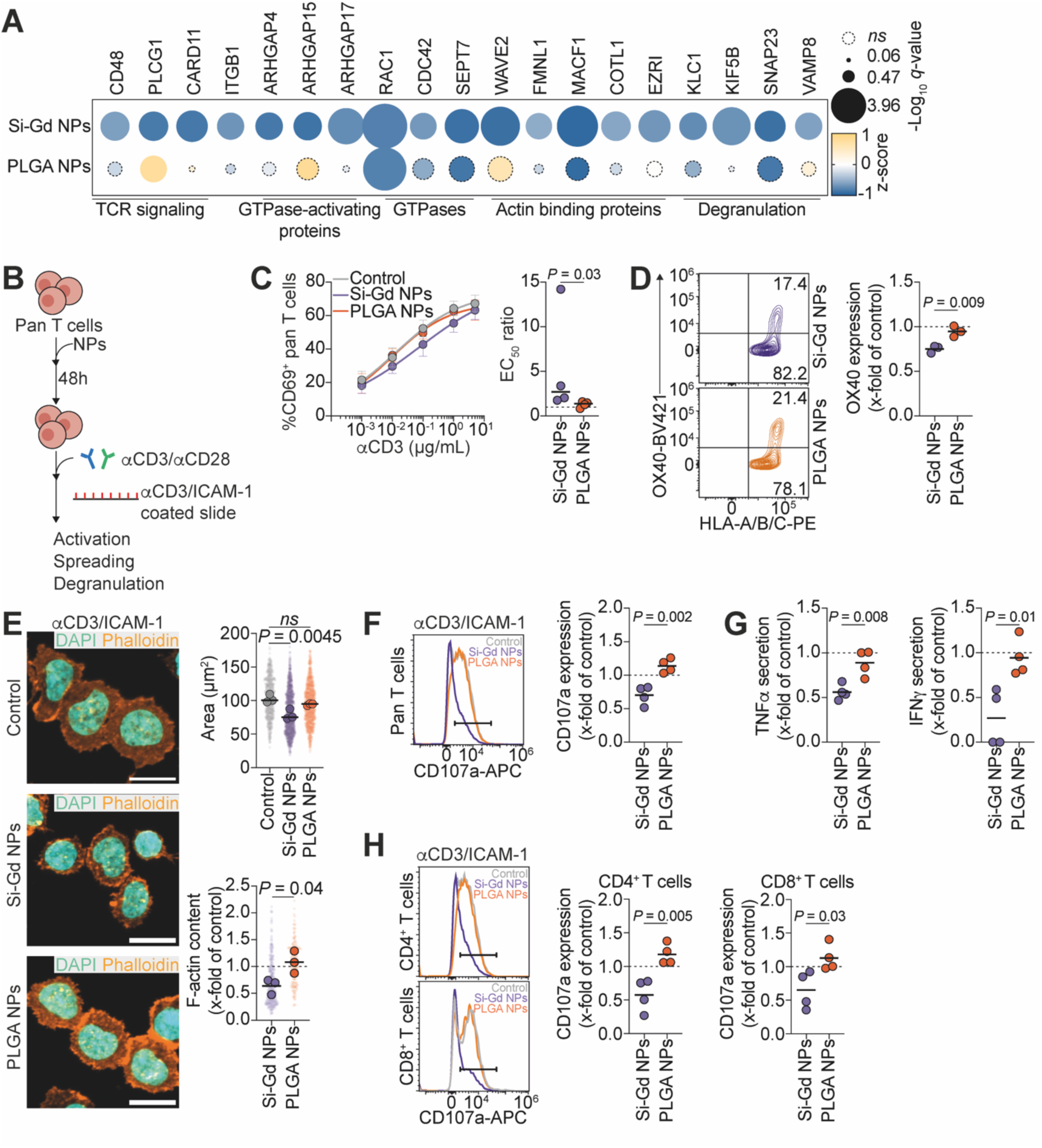
Silica-based gadolinium NPs treatment impairs pan T cells spreading and degranulation. **A**. Bubble plot showing Z-score values for proteins expression associated with pan T cells functions. Protein names are represented on the top while associations are represented on the bottom. Bubble size represents -Log_10_(*q*-value). Dashed lines bubbles correspond to non-significantly deregulated proteins. **B**. Schematic representation of the pipeline of the assessment of Si-Gd NPs and PLGA NPs functional impact on primary human pan T cells. **C**. Pan T cells sensibility to polyclonal activation. Left: Percentage of CD69-positive pan T cells assessed after culture on a culture plate coated with increasing concentration of anti-CD3 antibodies for 24h. Right: Half maximal effective concentration (EC_50_) calculated from the dose-response curve in the left graph. Data are representative of 4 independent experiments and analyzed by a Mann-Whitney test after assessment of their gaussian distribution by Shapiro-Wilk test. **D**. Flow-cytometry assessment of OX-40 expression at the pan T cells surface after activation with 5µg/mL of anti-CD3 and 5µg/mL of anti-CD28 antibodies for 24h. Left: Representative flow cytometry contour plots of the relative OX-40 expression in activated pan T cells previously treated for 48h with Si-Gd NPs (upper panel) or PLGA NPs (lower panel). Right: Quantification of OX-40 expression as fold change relative to untreated control. Data are representative of 3 independent experiments and analyzed by a Student’s t-test after assessment of their gaussian distribution by Shapiro-Wilk test. **E**. Nanomaterials-treated pan T cells spreading on an antigen-presenting cells mimicking surface. Left: Representative confocal micrographs of pan T cells spreading. In orange Phalloidin-iFluor488, in cyan nuclei (DAPI). Scale bar = 10µm. Right: Pan T cells median spreading area. Left: Relative filamentous actin content in pan T cells as fold change relative to untreated control. Data are representative of 3 independent experiments and analyzed by a One-way ANOVA test with original FDR method of Benjamini-Hochberg (Spreading area) or a Student’s t-test (F-actin content) after assessment of their gaussian distribution by Shapiro-Wilk test. **F**. Flow-cytometry analysis of pan T cells degranulation. Left: Representative histograms of CD107a (LAMP-1) expression at the pan T cells surface after activation with 5µg/mL of anti-CD3 and 5µg/mL of anti-CD28 antibodies for 4h. Right: Quantification of CD107a expression as fold change relative to untreated control. Data are representative of 4 independent experiments and analyzed by a Student’s t-test after assessment of their gaussian distribution by Shapiro-Wilk test. **G**. TNFa (left panel) and IFNg (right panel) secretion in the supernatant after activation with 5µg/mL of anti-CD3 and 5µg/mL of anti-CD28 antibodies for 4h. Data are represented as fold change relative to untreated control. Data are representative of 4 independent experiments and analyzed by a Student’s t-test after assessment of their gaussian distribution by Shapiro-Wilk test. **H**. Flow-cytometry analysis of CD4^+^ and CD8^+^ T cells degranulation. Left: Representative histograms of CD107a (LAMP-1) expression at the CD4^+^ T cells (upper panel) and CD8^+^ T cells (lower panel) surface after activation with 5µg/mL of anti-CD3 and 5µg/mL of anti-CD28 antibodies for 4h. Right: Quantification of CD107a expression as fold change relative to untreated control. Data are representative of 4 independent experiments and analyzed by a Student’s t-test after assessment of their gaussian distribution by Shapiro-Wilk test.

However, in the line with TCR signaling proteins downregulation observed in our proteomic data, Si-Gd NPs-treated pan T cells require higher anti-CD3/CD28 concentrations to reach similar activation levels than control and PLGA NPs-treated pan T cells, suggesting a reduced sensitivity to TCR activation (**Fig.4C**). We further confirm this weakened TCR signaling by measuring OX-40 expression, a co-stimulatory molecule selectively induced after TCR stimulation^[85,86]^. When normalizing to untreated pan T cells, we notice that Si-Gd NPs reduced by 25% OX-40 expression (**Fig.4D**).

Next, we sought to analyze NPs treatment effect on actin remodeling at the IS. We compare pan T cell spreading on glass surfaces coated with anti-CD3 together with a recombinant ICAM1-Fc chimera protein (**Fig.4B**). We monitor T cells morphology and cytoskeletal remodeling and observe that Si-Gd NPs, but not PLGA NPs, significantly reduce cell spreading and F-actin content (**Fig.4E**). T cells-mediated functions require not only spreading on the APCs surface but also degranulation. We thus assess the effect of NPs treatments on the degranulation of pan T cells as well as of their CD4^+^ and CD8^+^ subpopulations. To do so, we follow the cell surface expression of CD107a (LAMP-1) and cytokines release in T cells activated by anti-CD3/CD28 and ICAM1-Fc chimera protein. When compared to control, CD107a expression is significantly decreased by Si-Gd NPs treatment in pan T cells from all tested donors, hinting at an inhibition of their degranulation (**Fig.4F**). This phenotype is associated with a significant two-fold reduction in TNFα and IFNγ secretion (**Fig.4G**). We previously observe that CD4^+^ and CD8^+^ T cells internalized differentially the NPs (**Fig.1F**) and thus wonder whether the uptake level of NPs affect the T cells functionality. Hence, we further evaluate the CD107a expression of CD4^+^ and CD8^+^ T cells within the pan T cells population. Interestingly, CD4^+^ and CD8^+^ T cells exhibited similar significant two-fold diminution in CD107a expression upon Si-Gd NPs treatment, suggesting that their detrimental effect already occurs at low NP doses (**Fig.4H**).

Altogether, we highlight that Si-Gd NPs treatment widely impairs T cells activation, and functional responses to a polyclonal antigenic stimulation.

## Conclusion

Nanoimmunotherapies, combining nanotechnology and immunotherapy, represent a promising therapeutic strategy to improve anti-tumoral immune response. Understanding how NPs materials impact immune cells post-internalization is essential to ensure their safety profile and rationally guide the selection of specific NPs based on the therapeutic need.

Here, we present a pioneering analysis of the material impact from untargeted and drug-free NPs on two human primary immune cell types. Through a pre-screen combining cytotoxicity measurement and whole proteome analysis of a panel of five different nanomaterials under preclinical and clinical evaluation, we first highlight the toxicity of oxCNTs and terbium-based NPs (Tb and Si-Tb NPs) on NK and pan T cells. Overt toxicity at low concentration induced by oxCNTs might be partially explained by the downregulation of a wide range of proteins associated with cellular metabolic processes as well as ribosomal associated proteins, as observed by whole proteome analysis (**Fig.S3A**). Of note, ribosomal associated proteins modulation was previously observed following co-culture of Jurkat T cells with carbon-based NPs^[87]^.

Based on this pre-screen, we identify Si-Gd NPs and PLGA NPs as promising candidates for further preclinical development. We further examine Si-Gd NPs and PLGA NPs interactions with immune cells using a proteogenomic approach. Bulk RNA-sequencing reveal that PLGA NPs triggered a strong transcriptional priming towards activation in NK and pan T cells. We additionally validate by flow cytometry that drug-free PLGA NPs exposure was sufficient to activate NK and pan T cells, further confirming the transcriptional priming. Upstream regulators and aggregate expression analysis reveal that this priming is, in part, mediated by TNF-α via NFκB and IFN-γ pathways. Investigation of the expected protein-protein interactions network among upregulated genes in PLGA NPs-treated NK- and pan T cells (**Fig.S4C-D**) further underpin the complex interaction between TNF, NFκB, and IFN pathways. If we identify the signaling pathway involved in NK- and pan T cells activation by PLGA NPs, we still lack the precise identification of the first events and targets engaged during the immune cells-NPs interaction. Hence, further evaluation to decipher to exact mechanism of NK and pan T cells activation are required.

More detailed investigations of NK functions upon NPs exposure reveal that PLGA NPs treatment efficiently enhance NK cells tumoricidal activity in cytokines-deprived environment. In depth analysis showcase that the NK cells functional enhancement is mediated by a better mature perforin polarization at the IS associated with a surge in cytotoxic cytokines release. Increased perforin polarization might be partially explained by an enhanced NK cells maturation and vesicles transport as observed in our proteogenomic data. In line with our results, another study recently demonstrated that drug-free polymer micropatches were able to activate murine neutrophils and induce an N1 anti-tumoral response^[88]^. Yet, our *in vitro* analysis does not allow to evaluate how NK cells priming integrate into the global adaptative immune system. Since NK cells are key player in the regulation of adaptive immune responses^[89]^, notably by the recruitment dendritic cells via XCL1^[40]^ (that is overexpressed by PLGA NPs-treated NK cells), thorough *in vivo* evaluation remained to be performed to decipher *ex vivo* primed NK cells tumoricidal enhancement on tumor burden regression and systemic immune response induction.

In parallel, explorations of pan T functions expose that Si-Gd NPs treatment widely impaired their activation, and functional responses to a polyclonal antigenic stimulation. Detailed analysis reveals that impairment was similar in CD4^+^ and CD8^+^ T cells despite different internalization rates, suggesting that their detrimental effect already occurs at low doses. Si-Gd NPs are under clinical evaluation as a MRI contrast agent^[90]^ and radio-enhancer^[91]^ (NCT04789486). Si-Gd NPs increase the radiation effects on tumor cells, inducing higher immunogenic cell death levels and higher immune cells recruitment to the tumor bed^[92]^. Therefore, a careful assessment of the impact of Si-Gd NPs on recruited immune cells is necessary. In addition, Si-Gd NPs were recently targeted with a programmed death-ligand 1 (PD-L1) VHH to monitor immune checkpoint molecules expression *in vivo* through medical imaging^[30]^. However, human effector T cells are also PD-L1^[93,94]^. These results suggest that Si-Gd NPs development as medical imaging tracers to assess immune infiltration in tumors and to predict response to immunotherapy treatments should be carefully evaluated. Interestingly, a previous study exhibit that ultrasmall silica NPs were able to induce a dose-dependent CD4^+^ and CD8^+^ T cells activation^[34]^. These ultrasmall silica NPs possess a hydrodynamic diameter similar than our Si-Gd NPs, suggesting than metallic loading can impact T cells response. Further investigations in that sense should be performed to evaluate other metals (such as gold or silver NPs). While transcriptomic analysis reveals that PLGA NPs seemed to promote a T_H_1 anti-tumoral polarization signature, working with healthy pan T cells we were not able to investigate this aspect of the NPs impact. Indeed, healthy pan T cells do not possess a tumor antigen-specific TCR. Further evaluation working with T cells sourced from cancer patients or TCR-engineered T cells are required to evaluate the function impact of the transcriptional T_H_1 polarization.

Our findings provide new insights for rational selection of NPs materials in nanoimmunotherapeutic approach. While additional *in vivo* investigations are required, we identify drug-free PLGA NPs as suitable and promising candidates for further targeting approaches aiming to reactivate the immune system of cancer patients. Our results suggest that intrinsic PLGAs NPs materials impact could synergize with immuno-activating agents to efficiently target and modulate the immune system.

## Experimental section

### Nanoparticles synthesis

Nanoparticles were synthetized as previously described. Briefly, carbon nanotubes have been shortened under strong acid conditions (H_2_SO_4_/HNO_3_ 3:1) and sonication for 24h to generate a high amount of carboxylic groups at the tips and around the side walls^[95]^. The ultra-small metal-based NPs are composed by a polysiloxane core, and a shell comprising amine function and metal complexed by DOTAGA (in same amount) and were provided by the Institut Lumière-Matière of the University of Lyon. Si-Gd were synthetized by a top-down process^[31]^ and Si-Tb were synthetized in a one-pot protocol^[32]^. Lanthanide-doped La_0.9_Tb_0.1_F_3_ NPs (hereafter, Tb NPs) were synthetized by dropwise addition of 0.9 LaCl_3_ and 0.1 TbCl_3_ to 3 NH_4_F followed by heating at 150°C for 12 mins and purification, as described in^[96]^. Finaly, hybrid PLGA-lipid NPs were synthetized via self-assembly of poly(D,L-lactide-*co*-glycolide) acid (30-60 kDa, lactide:glycolide 50:50; Sigma #P2191) and 1,2-distearoyl-sn-glycero-3-phosphoethanolamine-N-[carboxy(polyethylene glycol)-2000] (sodium salt) (DSPE-PEG-CO₂H; Avanti Polar Lipids #880135) through a one-step nanoprecipitation method as described previously in^[97]^. Briefly, PLGA polymer was dissolved with or without 0.2% Cyanine 5.5 carboxylic acid (Lumiprobe #17090,; λ_ex_/λ_em_: 684nm/710 nm) in acetonitrile at a concentration of 5 mg/mL. DSPE-PEG-CO₂H, at a weight ratio of 20% relative to PLGA polymer, was dissolved in 10 mL of 4 wt% ethanol aqueous solution and stirred vigorously at 65 °C. The PLGA solution was then added dropwise into the lipid solution using a syringe pump (0.5 mL/h) under constant stirring. The entire mixture was kept under gentle stirring for 2 hours at room temperature under a chemical hood. The remaining organic solvent and unloaded molecules were removed by washing the NP solution with ultrapure water using an Amicon Ultra-15 centrifugal filter from Millipore, France (cut-off: 50 kDa; 3 cycles, 3,000*g*, 10 min) via tangential centrifugation. The final NP formulation was reconstituted in 1 mL of ultrapure water.

### Dynamic Light Scattering (DLS)

DLS measurements were conducted using a nano-ZS instrument (Malvern). The suspensions of NPs were prepared in a solution of nanopure water (Milli-Q). DLS measurements were performed in sets of 10 acquisitions. The average hydrodynamic diameters of the NPs were determined by analyzing the DLS correlation function through a regularization fitting method.

### Human NK and pan T cells isolation

Peripheral blood mononuclear cells (PBMC) were obtained by density gradient centrifugation (1,200*g*, 20mins) of buffy coats from healthy volunteer blood donors under written informed consent recruited at Établissement Français du Sang Grand-Est, Strasbourg, France (agreement A122395/2022). Following erythrolysis (MiltenyiBiotec #130-094-183), NK cells and pan T cells were isolated from PBMC by negative magnetic cell-sorting (MiltenyiBiotec #130-092-657and #130-096-535) following manufacturer instructions. The cell purity was then controlled by flow cytometry on a Attune NxT (Invitrogen) flow cytometer using PE anti-CD56 (1:100, MiltenyiBiotec # 130-113-312) and FITC anti-CD3 (1:100, MiltenyiBiotec #130-126-882) staining and ranged from 90% to 99.2% (median 96.3%). Gating strategy is shown in Figure S2.

NK- and pan T cells (10^6^ cells/mL) were cultured for further experiment in R10 medium: RPMI 1640 (Gibco #72400054) containing 2mM Glutamine, 25mM HEPES and completed with 10 % v/v of (fetal bovine serum) FBS (Gibco), 100 U/mL penicillin, 100 μg/mL streptomycin (PanBiotech #P06-07100) and 50 μM β-mercaptoethanol (Gibco #11508916).

### Cell lines and cell line engineering

The human chronic myelogenous leukemia cell line K562 (ATCC CCL-243) was cultured under standard conditions (37°C, 5% CO_2_) using R10 medium. Cell viability *in vitro* was assayed before functional experiment by a Countess 3^®^ automated cell counter (ThermoFisher).

For immune synapse formation, K562 cells were engineered to express a palmitoylated tdTomato. Briefly, the tdTomatoDNA fragment from the Addgene plasmid tdTomato-Lifeact-7 (#54528) was amplified by PCR to add the palmitoylation sequence of the *GAP43* gene and then cloned in pJET 1.2 vector according to manufactory instructions (ThermoFisher #K1231). The generated Mb-tdTomato fragment was then cloned in a lentivirus pLSFFV-IRES-Blasticidin vector. Lentivirus generated from the pLSFFV-Mb-tdTomato-IRES-Blasticidin construct were produced by transfection together with 3 additional vectors (pLP1, pLP2 and pLP3-VSV plasmids) in HEK293T cells using JetPRIME transfection reagent (Polyplus).

For K562 cells transduction, 6 wells plate was coated with Retronectin (Takara, 10µg/cm^2^) for 2h at room temperature. Wells were washed in PBS before blocking with 2% BSA in PBS for 30 min. After a wash with PBS, 400.000 K562 cells were seeded and let adhere overnight under standard conditions (37°C, 5% CO_2_). The day after, lentiviruses encoding for the pLSFFV-Mb-tdTomato-IRES-Blasticidin construct were added in the presence of polybrene (10µg/mL). After one day of transduction, selection with blasticidin (5µg/mL) was performed until highly fluorescent cells were FACS sorted.

### Nanoparticles impact on immune cells viability

Negatively sorted NK and pan T cells (100,000/wells) were treated with increasing concentrations of each NPs types in R10 medium. Cells were treated for 48h under standard conditions (37°C, 5% CO_2_). Immune cells viability was measured at the endpoint using the CellTiterGlo^®^ luminescent cell viability assay (Promega #G7570).

### Immune cells RNA extraction and bulk RNA-sequencing

#### RNA extraction and quantification

After 48h of culture, NK and pan T cells untreated or treated with NPs at the indicated concentrations were harvested and washed in ice-cold PBS. Total RNA was isolated using the RNAeasy kit (Qiagen #74136) with on-column DNAse I digestion (Qiagen #79256) according to manufacturer’s instruction. RNA was eluted in 50μL final volume, and its concentration was assessed with the NanoPhotometer^®^ N60 (Implemen). RNA integrity was assessed using a total RNA Pico Kit by Bioanalyzer 2100 Instrument (Agilent Technologies). All samples had RNA integrity numbers above 7.

#### Library preparation and sequencing

Sequencing libraries were prepared using “NEBNext Ultra II Directional RNA Library Prep Kit for Illumina” and enriched in mRNA using “NEB Ultra II polyA m RNA magnetic isolation” kit (New England Biolabs). Libraries were pooled and sequenced (single-end, 100bp) on a NextSeq2000 according to the manufacturer’s instructions (Illumina Inc.).

#### Multidimensional scaling (MDS) and differential expression analysis

For each sample, quality control was carried out and assessed with the NGS Core Tools FastQC^[98]^. Sequence reads (minimum 22.4 million) were mapped to *Homo sapiens* hg19 using STAR^[99]^ to obtain a BAM (Binary Alignment Map) file. An abundance matrix was generated based on read counts identified by HTSeq-count^[100]^. Multidimensional scaling (MDS) was performed on gene expression counts normalized using the DESeq2 R package^[101]^ to investigate relative similarities across the different conditions. The first two principal components were plotted against each other, along with their respective variances explained. Differential expression analysis was then performed between the NPs treated conditions and the control using DESeq2^[101]^ package of the Bioconductor framework for RNASeq data^[102]^. Volcano plots were constructed based on log_2_ fold change (≥ or ≤ 2) and −log_10_ FDR adjusted *p*-value (<1.3).

#### Erichment analysis

Enrichment analysis of Gene Ontology (GO) terms a was conducted using Metascape (http://metascape.org)^[103]^. Functional analysis of gene expression changes was undertaken using Ingenuity Pathway Analysis (IPA, Ingenuity Systems).

#### Estimation of aggregate expression

Genes related to TNFα signaling via NFκB and IFNγ pathways were curated from the MSigDb Hallmark 2024 database (**Table S4**). For each experimental condition, aggregate expression levels of the indicated gene signatures were estimated by first performing a normalization on the cell counts. The normalized counts were then z-scored by gene (across all the conditions), after which the genes of interest were subsetted and their distribution of z-scored gene counts visualized as violin plots.

### RT-qPCR analysis

Downstreamed regulators expression was performed by generating complementary DNA (cDNA) with the High-Capacity cDNA Reverse Transcription Kit (Applied Biosystems #4368814), according to the manufacturer’s instructions. qPCR analysis on each biological sample was performed using technical replicates with the TaqMan system on a QuantStudio™ 3. The cDNA concentration of target genes was normalized by amplification of GAPDH and fold changes in gene expression were obtained using the 2^-ΔΔCt^ method. TaqMan probes were NFKB1 (Hs00765730_m1), NFKB2 (Hs01028890_g1), RELA (Hs01042014_m1), RELB (Hs00232399_m1), IRF1 (Hs00971965_m1), GAPDH (Hs02786624_g1).

### Whole proteome analysis

#### Protein extraction and samples preparation

After 48h of culture, NK and pan T cells untreated or treated with NPs at the indicated concentrations were harvested and washed in ice-cold PBS. Immune cells dry pellets were flash frozen in liquid nitrogen. Cellular pellets were resuspended in 2% SDS, 62.5 mM Tris-HCl pH = 6.8 and lysed using a water bath sonicator cooled with ice. Protein concentration was estimated using the Biorad DC kit (Hercules). Proteins (2µg) were prepared using a modified SP3 workflow based on^[104]^. Briefly, proteins were reduced 30 minutes at 37°C with dithiothreitol (final conc. 12mM) and alkylated 30 minutes, RT, in the dark with iodoacetamide (final conc. 40mM). SP3 magnetic beads (Sera-Mag SpeedBeads) were rinsed 3 times with H_2_O before being added to the sample (ratio 1:10 protein/beads). Acetonitrile (final conc. 50% v/v) was added to precipitate the proteins on the beads and the samples were incubated for 15min, RT, with agitation. The beads were washed twice with 200μL of 80% ethanol and once with 180μL of acetonitrile before being resuspended in 40μL of ammonium bicarbonate (100mM) followed by 5min sonication in a water bath. Trypsin/Lys-C was added to achieve a final ratio of 1:10 (enzyme:protein) and the proteins were digested overnight, 37°C, 600rpm. Samples were acidified with formic acid to a final conc. of 1% v/v and centrifuged for 10min at 3500rpm. The samples were incubated for 10 minutes on the magnetic rack and the supernatants containing the peptides were transferred to a new plate. Peptide clean-up was performed on a Bravo AssayMap (Agilent) using 5μL RP-C18 cartridges (Agilent) following the manufacturer’s instructions.

#### nLC-MS/MS analysis

After evaporation, peptides were resuspended in H_2_O/ACN/FA (98/2/0.1) and 1/6th of the peptides were injected in randomised order on a nanoAcquity (Waters) - Q-Exactive HF-X coupling (Thermo Fisher Scientific). Peptides were separated using a 79min gradient at a flow rate of 400nL/min. The amount of solvent B (ACN/FA, 99.9/0.1) started at 1%, increased to 8% in 2min and then to 35% B in 77min. The column was washed by increasing the percentage of B to 90% in 1min and for 5min before decreasing to 1% B in 2min and for 2min to re-equilibrate the column. MS analysis was performed using a TOP20 data-dependent acquisition. The scan range was 375 to 1500m/z with a dynamic exclusion of 40s. For precursor analysis, a resolution of 120,000 was used with an AGC target of 3.0E+06 and a maximum injection time of 60ms. For fragment analysis, the resolution was 15,000 with an AGC target of 1.0E+05, a maximum injection time of 60ms and an isolation window of 2m/z.

#### Data treatment and differential analysis

Data searches were performed on a local Mascot server (Matrix Science) using a database containing all human protein entries from the UniProtKB/SwissProt database and classical MS contaminant proteins. A tolerance of 5 ppm for precursors and 0.05 Da for fragments was applied. Carbamidomethylation of cysteine residues was defined as a fixed modification, while acetylation of the N-termini of the proteins and oxidation of methionines were defined as variable modifications. Proline studio^[105]^ was used for validation of protein identifications and quantification using a 1% FDR at both the protein and PSM levels. The Prostar software (v 1.22.6)^[106]^ was used for the differential analysis. The filtering keeps only proteins with at least two values for one condition. The abundance was normalised using a quantile centering normalisation over all the analysis. The imputation of the Partially Observed Value (POV) was realised using Structured Least Square Adaptative (SLSA) imputation whereas the imputation of the values Missing in an Entire Condition (MEC) was realised using det quantile imputation. The hypothesis testing used a Limma test for one condition in comparison to the control. Finally, the P-value calibration was realised using Benjamini-Hochberg calibration. Results were filtered to obtain a FDR of around 1%.

### NK cells activation and K562 killing assay

For NK cells activation, 40,000 K562 target cells were seeded per wells in a U-bottom 96-wells plate. Untreated or NPs-treated NK cells were washed and resuspended at 4.10^6^ cells/mL in R10 medium and added at increasing effector-to-target ratios (0:1: 0.625:1; 1.25:1; 2.5:1 and 5:1) to K562 cells. NK were co-incubated with their target cells for 4h at 37°C, 5% CO_2_. After incubation, the cells were washed in PBS and non-viable cells were stained with Fixable Viability Dye-eFluor450 (eBioscience #65-0863-14) for 15 minutes at room temperature in the dark. Aspecific antibodies binding was minimized by treating the cells with TruStain FcX anti-CD32/CD16 blocking antibody (1:50, BioLegend # 422301) for 20 minutes at 4°C. Then, cells were stained with PE anti-CD56 (1:100, MiltenyiBiotec #130-113-312) and PE-Vio700 anti-CD69 (1:50, MiltenyiBiotec #130-112-615) and APC anti-NKG2D (1:50, MiltenyiBiotec #130-111-846) for 15 min at 4°C. Samples were acquired with Attune NxT (Invitrogen) flow cytometer and data were analyzed using FlowJo™ v10 Software (ThreeStar). Gating strategy is shown in Figure S6A-B.

### NK cells immune synapse formation

Conjugates between untreated or NPs-treated NK cells and K562-palmitoylated tdTomato target cells at a 2:1 ratio were formed in suspension for 20 min at 37°C in serum-free R10 medium. Cells were then gently mixed and transferred to 0.01% poly-L-lysine coated 12-wells ibidi slides (ibidi #81201). Slides were incubated for 25 minutes at 37°C/5% CO_2_ and then fixed with 2% PFA in PBS for 20 min at room temperature. Fixative was removed and wells were rinsed three times with 150 μl PBS. Cells were permeabilized with 0.1% Triton X-100 / 2% BSA in 1x PBS for 5 min and unspecific antibody binding was blocked for 1 h in blocking solution (3% bovine serum albumin-BSA, 5% FBS, 0.01% Triton X-100, in PBS). Immunostaining was performed with purified mouse anti-perforin (10µg/mL, BioLegend, clone dG9), incubated overnight at 4°C followed by secondary antibodies (Goat anti-mouse-AlexaFluor 647, 1:250, ThermoFisher #A-21236) and Phalloidin-iFluor488 (1:1,000, Abcam #ab176753) for 1h room temperature prior to slides mounting (Fluoromount/DAPI). Background and nonspecific staining controls were used.

Perforin-stained NK-K562 conjugates were imaged with a 60X water-immersion objective on an inverted OIympus Spinning-disk, and *z*-series images were acquired with a space of 0.45 μm. Images were processed using ImageJ software (National Institutes of Health). For scoring of cytotoxic granules distance to the immune synapse (based on perforin staining), 20 conjugates between NK cells and K562 target cells were chosen randomly per condition. Distance quantification was performed using the ImageJ macro “Shortest_distance_between_objects” as described in^[107]^.

### Pan T cells activation analysis

For T cells activation, F-bottom 96-wells plate were coated increasing anti-CD3 concentrations (ranging from 0.001 to 10μg/mL, BioLegend, clone OKT3) overnight at 4°C. Wells were washed once in PBS before adding the cells. Untreated or NPs-treated pan T cells were washed and resuspended at 10^6^ cells/mL in R10 medium and anti-CD28 (5μg/mL, BioLegend, clone 28.2) was added to each condition. 100,000 cells per wells were incubated for 24h at 37°C, 5% CO_2_. After incubation, the cells were washed in PBS and non-viable cells were stained with Fixable Viability Dye-eFluor780 (eBioscience #65-0865-14) for 15 minutes at room temperature in the dark. Aspecific antibodies binding was minimized by treating the cells with TruStain FcX anti-CD32/CD16 blocking antibody (1:50, BioLegend # 422301) for 20 minutes at 4°C. Then, cells were stained with FITC anti-CD3 (1:100, MiltenyiBiotec #130-126-882), PE-Vio700 anti-CD69 (1:50, MiltenyiBiotec #130-112-615) and Brilliant Violet 421 anti-OX40 (1:20, BioLegend, clone Ber-ACT35) for 15 min at 4°C. Samples were acquired with Attune NxT (Invitrogen) flow cytometer and data were analyzed using FlowJo™ v10 Software (ThreeStar).

### Pan T cells spreading and F-actin analysis

For T cells spreading, 8-wells ibidi slides (ibidi #80841) were coated with 0.01% poly-L-lysine for 45 min at room temperature. After washing in PBS, the coverslips were coated with anti-CD3 (10μg/mL, BioLegend, clone OKT3) and rhICAM1-Fc (2μg/mL, BioLegend #552906) overnight at 4°C. Coverslips were washed once in PBS before adding the cells. Untreated or NPs-treated pan T cells were washed and resuspended at 10^6^ cells/mL in serum-free R10 medium for 1h at 37°C/5%CO_2_. After, 100,000 cells per wells were incubated with anti-CD28 (5μg/mL, BioLegend, clone 28.2) for 25 min at 37°C, 5% CO_2_. Coverslips were then washed once with PBS and fixed with % PFA in PBS for 20 min at room temperature. Fixative was removed and wells were rinsed three times with 150 μl PBS. Cells were permeabilized with 0.1% Triton X-100 / 2% BSA in 1x PBS for 5 min and F-actin was stained by Phalloidin-iFluor488 (1:1,000, Abcam #ab176753) for 1h room temperature prior to slides mounting (Fluoromount/DAPI).

Phalloidin-stained pan T cells were imaged with a 60X water-immersion objective on an inverted OIympus Spinning-disk, and *z*-series images were acquired with a space of 0.45 μm. Images were processed using ImageJ software (National Institutes of Health). To analyze cell spreading on surfaces, only the z-stack plane corresponding to the contact between cells and the surface was considered, and a projection of the DAPI stain was used to individualize each cell. Briefly, cell areas were obtained by polygon selection definition and spreading area as well as F-actin intensity at the immunological synapse were measured. F-actin intensity was normalized by each cell area.

### Pan T cells degranulation analysis

For T cells degranulation, F-bottom 96-wells plate were coated anti-CD3 (10μg/mL, BioLegend, clone OKT3) and rhICAM1-Fc (2μg/mL, BioLegend #552906) overnight at 4°C. Wells were washed once in PBS before adding the cells. Untreated or NPs-treated pan T cells were washed and resuspended at 10^6^ cells/mL in R10 medium and anti-CD28 (2μg/mL, BioLegend, clone 28.2) was added to each condition. 100,000 cells per wells were incubated for 1h at 37°C, 5% CO_2_ in presence of APC-conjugated anti-CD107a (1:50, MiltenyiBiotec #130-111-847). After 1h, protein transport inhibitor cocktail (1:500, eBioscience) was added in each well and cell were incubated for addition 4h at 37°C, 5% CO_2_. After incubation, the cells were washed in PBS and non-viable cells were stained with Fixable Viability Dye-eFluor780 (eBioscience #65-0865-14) for 15 minutes at room temperature in the dark. Aspecific antibodies binding was minimized by treating the cells with TruStain FcX anti-CD32/CD16 blocking antibody (1:50, BioLegend # 422301) for 20 minutes at 4°C. Then, cells were stained with VioBright B515 anti-CD3 (1:50, MiltenyiBiotec #130-126-882), Pacific Blue anti-CD4 (1:100, BioLegend, clone RPA-T4) and PE anti-CD8 (1:100, BioLegend, clone RPA-T8) for 15 min at 4°C. Samples were acquired with Attune NxT (Invitrogen) flow cytometer and data were analyzed using FlowJo™ v10 Software (ThreeStar).

### Cytokine measurement

Supernatants from 96-well plates were aliquoted after 4h of NK-K562 cells co-incubated or after 24h of pan T cells activation and stored at −20 °C. Cytokines were quantified by custom LEGENDplex (5-plex, BioLegend # 740510) with the Attune NxT Flow Cytometer. Cytokine assays were analyzed using the LEGENDplex software Qognit (https://legendplex.qognit.com).

### Statistics

Statistical analysis was performed with GraphPad Prism 9.5. The normal distribution of the data sets was assessed by the Shapiro–Wilk normality test. According to the number of data sets compared, Student’s t-test test or One-way ANOVA test with original FDR method of Benjamini-Hochberg were applied (*p<0.05; **p<0.01, ***p<0.001, ****p<0.0001). In all cases, the α-level was set at 0.05. All the data in graphs were presented as median ± standard deviation.

## Data Availability

Raw RNA-seq data have been deposited in the EMBL-EBI ArrayExpress archive (accession number E-MTAB-14550). Complete proteomics dataset has been deposited to the ProteomeXchange Consortium via the PRIDE partner repository^[108]^ (accession number PXD056695). All the other data are available within the article and its Supplementary Information. Raw data are accessible through reasonable requests to the corresponding authors.

## Acknowledgements

We thank all members of the JGG and AD labs for helpful discussions. The authors wish to thank R. Soltani and A. Bianco for kindly providing the oxidized carbon nanotubes. We thank Pascal Kessler (PICSTRA, CRBS) for assistance in imaging as well as Claudine Ebel and Muriel Philipps (IGBMC, Strasbourg, France) for assistance in FACS sorting. We are grateful to Alexandre F. Carisey (St. Jude Children’s Research Hospital, Memphis, USA) for sharing K562 cells. This work was supported by a fellowship from the French Ministry of Science (MESRI) and a fourth-year thesis fellowship from the Fondation ARC pour la recherche sur le cancer to VM. Work and people in the lab of JGG are supported by the INSERM, the University of Strasbourg, as well as by La Ligue Nationale Contre le Cancer (LNCC) and the Association pour la Recherche contre le Cancer (ARC). LB is funded by FRM (Fondation pour la Recherche Médicale). We are also thankful for recent donators (Rohan Athlétisme Saverne and Traileurs de la Rose) to support our work. AD acknowledges funding from the Cancéropole Grand Est, the Fondation Française contre le Myélome et les Gammapathies, the International Myeloma Foundation, as well as support from the Institut de Cancérologie Strasbourg Europe and the European Research Council (ERC) under the European Union’s Horizon 2020 research and innovation program (ERC Starting Grant TheranoImmuno, grant agreement No. 950101) and the Ligue Nationale Contre le Cancer. Additionally, this work was also supported by the CNRS, the University of Strasbourg, the Agence National de la Recherche, the French Proteomics Infrastructure (ProFI FR2048; ANR-10-INBS-08-03), the Interdisciplinary Thematic Institute IMS, the drug discovery and development institute, as part of the ITI 2021-2028 program of the University of Strasbourg, CNRS and Inserm supported by IdEx Unistra (ANR-10-IDEX-0002) and SFRI-STRAT’US project (ANR-20-SFRI-0012) under the framework of the French Investments for the Future Program. We finally also acknowledge funding by IBiSA and Region Grand Est.

## Disclosure and competing interest statement

A.D., O.T., and F.L. are shareholders of NH Theraguix who is translating to the clinic Gd-NPs. The other authors declare that they have no conflict of interest.

## Authorship contributions

Conceptualization, V.M., O.L., J.G.G., and A.D.; Methodology, V.M., and O.L.; Investigation, V.M., M.C.D., J.B., A.L., A.P., and C.M.; Resources, M.B., S.G., L.J.C., F.L., and O.T.; Formal Analysis, V.M., O.L., A.D., M.C.D., L.B., S.H.,, A.P., T.S., A.M., M.R., C.C., and R.C.; Writing – Original Draft, V.M., O.L., J.G.G., and A.D.; Writing – Review & Editing, V.M., O.L., C.C., R.C., L.J.C., F.L., O.T., J.G.G., and A.D.; Supervision, C.C., R.C., J.G.G., and A.D.; Funding Acquisition, J.G.G., and A.D.

---

**Supplementary Figure 1 related to Figure 1.**
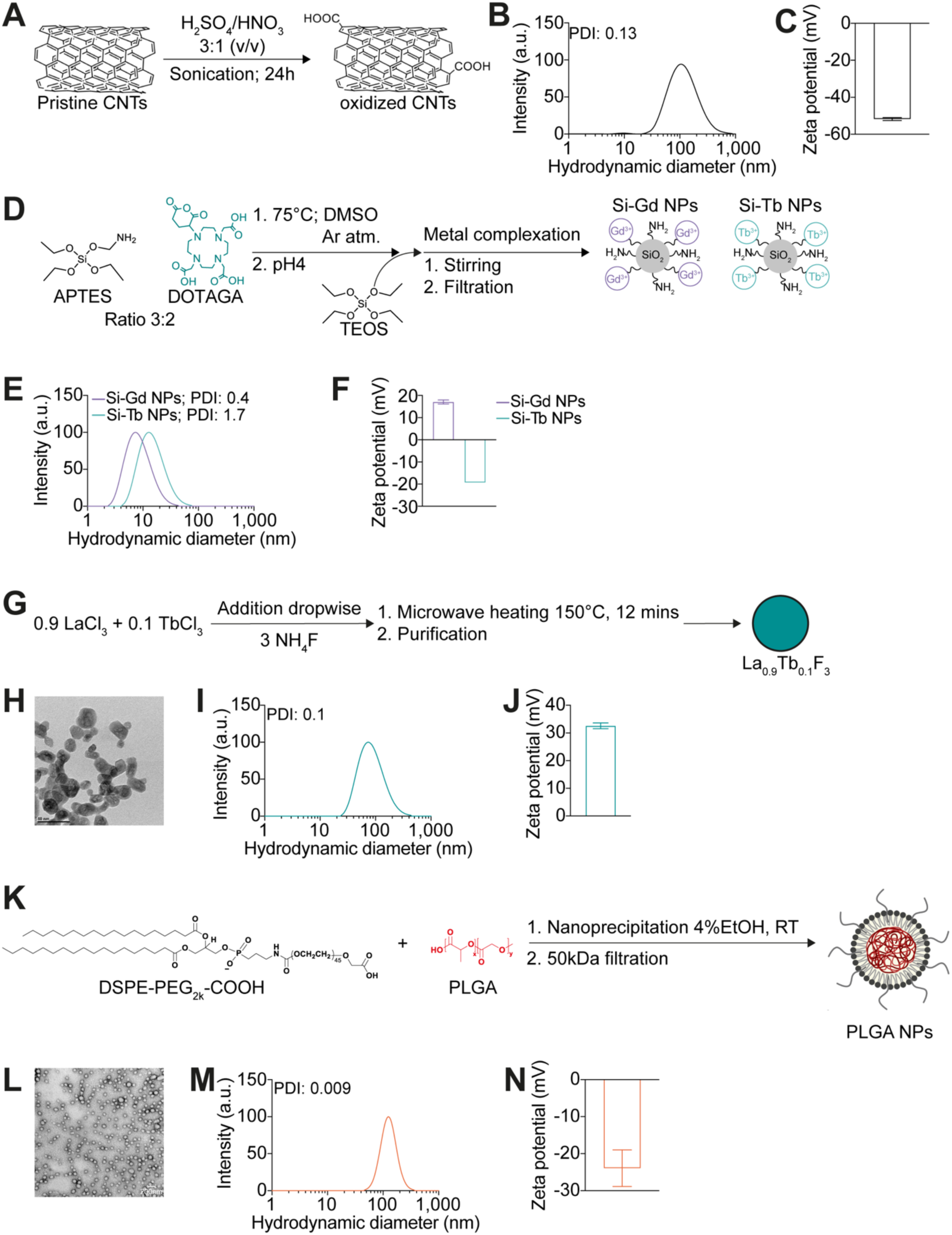
Nanomaterials synthesis and characterization. **A.** Prisitin CNTs were shortened under strong acid conditions (H_2_SO_4_/HNO_3_ 3:1) and sonication for 24h to generate a high amount of carboxylic groups. **B-C.** oxidated CNTs size and charge were measured by DLS. **D.** Ultrasmall polysiloxane-based gadolinium (Si-Gd) NPs were synthesized by a top-down method from core (gadolinium oxide) shell (polysiloxane) NPs whereas ultrasmall polysiloxane-based terbium (Si-Tb) NPs were synthesized with a bottom-up one pot synthesis. **E-F.** Si-Gd NPs and Si-Tb NPs size and charge were measured by DLS. **G.** Terbium fluoride (Tb) NPs were synthetized by dropwise addition of 0.9 LaCl_3_ and 0.1 TbCl_3_ to 3 NH_4_F followed by heating at 150°C for 12 mins and purification. **G.** Transmission electron microscopy of Tb NPs. **I-J.** Tb NPs size and charge were measured by DLS. **K.** PLGA NPs were synthetized by self-assembly of DSPE-PEG_2k_-COOH and PLGA through a one-step nanoprecipitation. **L.** Transmission electron microscopy of PLGA NPs. **M-N.** PLGA NPs size and charge were measured by DLS.

**Supplementary Figure 2 related to Figure 1.**
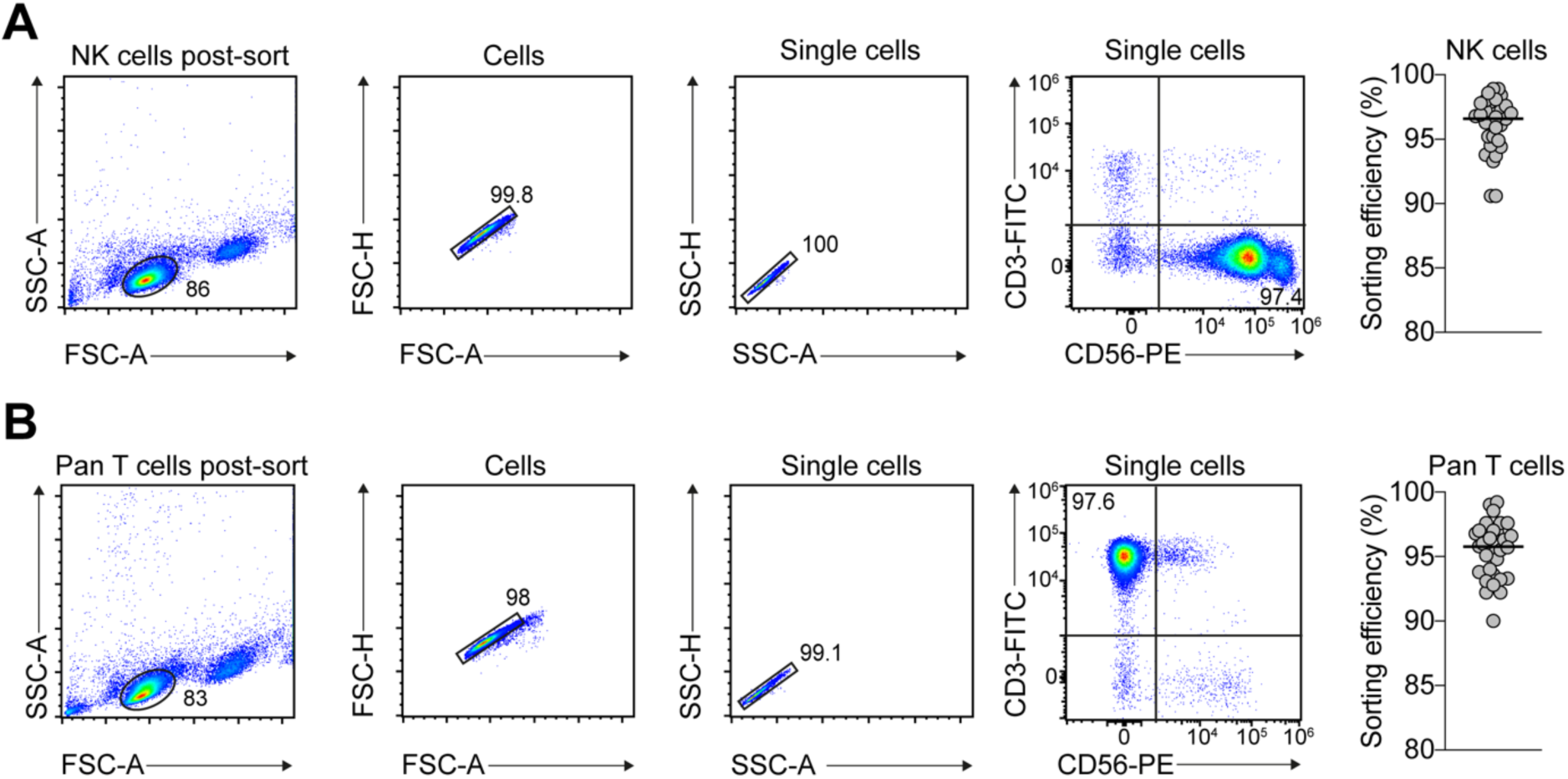
NK cells and pan T cells isolation. NK cells and pan T cells immunophenotyping. **A.** Left: Gating strategy for assessment of NK cells sorting efficiency. Right: Percentage of pure CD3^-^ CD56^+^ NK cells after sorting. Data are representative of 31 independent experiments. Median of the data is presented. **B.** Left: Gating strategy for assessment of pan T cells sorting efficiency. Right: Percentage of pure CD3^+^ pan T cells after sorting. Data are representative of 30 independent experiments. Median of the data is presented.

**Supplementary Figure 3 related to Figure 1.**
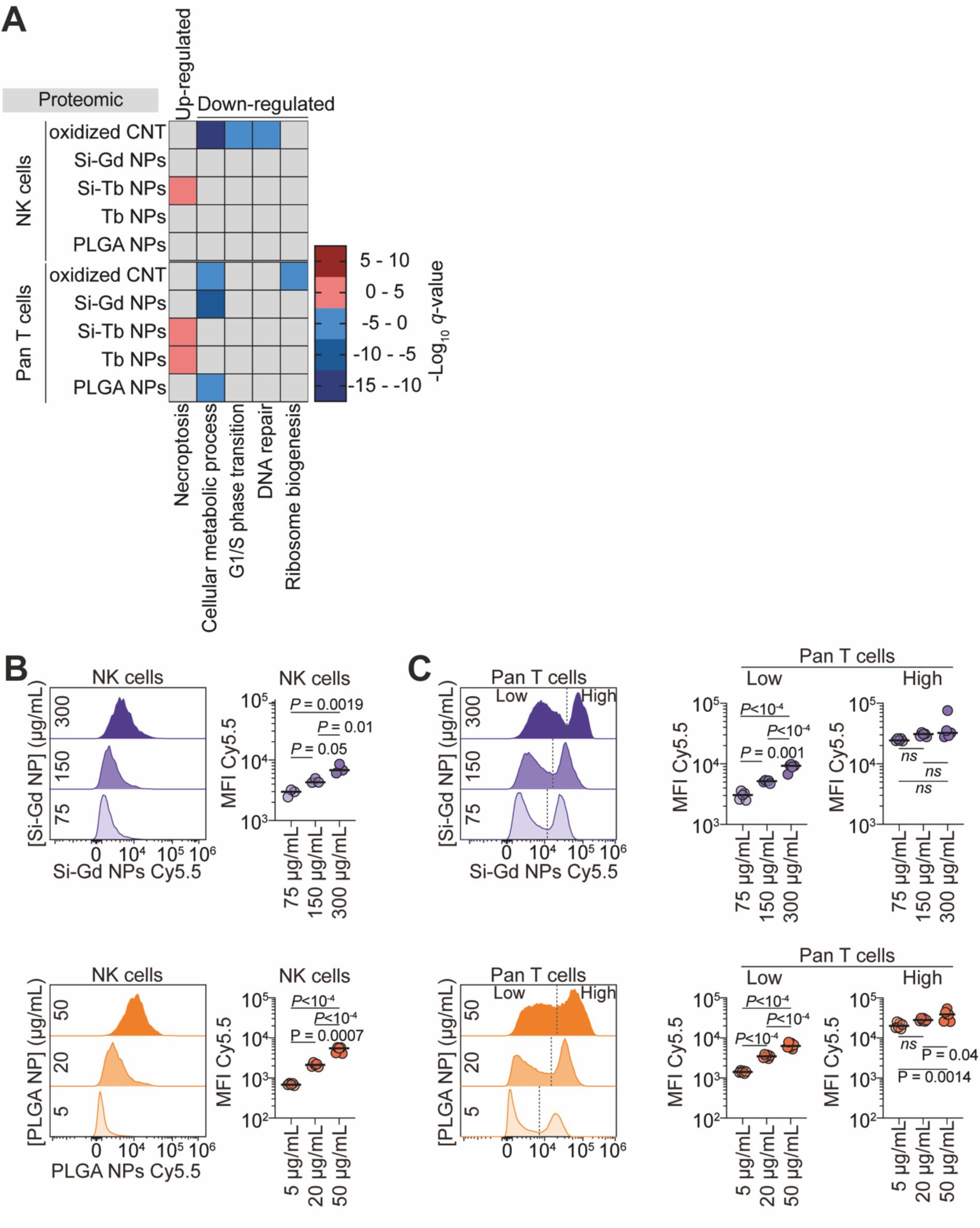
**A.** Heat-map showing -Log_10_(*q*-value) for up-regulated and down-regulated gene ontology terms in NK cells and pan T cells upon 48h treatment with different nanomaterials. **B.** Flow-cytometry assessment of nanomaterials internalization dose-dependency in NK cells after 48h co-incubation at 37°C. Left: Representative histograms of the relative internalization of Si-Gd NPs (upper panel) and PLGA NPs (lower panel). Right: Quantification of the mean fluorescent intensity signal of the Cyanine5.5-labelled nanomaterials. Data are representative of 3 to 6 independent experiments and analyzed by a One-way ANOVA test with original FDR method of Benjamini-Hochberg after assessment of their gaussian distribution by Shapiro-Wilk test. **C.** Flow-cytometry assessment of nanomaterials internalization dose-dependency in pan T cells after 48h co-incubation at 37°C. Left: Representative histograms of the relative internalization of Si-Gd NPs (upper panel) and PLGA NPs (lower panel). Right: Quantification of the mean fluorescent intensity signal of the Cyanine5.5-labelled nanomaterials in low and high internalizing population as gated on the histograms. Data are representative of 5 to 6 independent experiments and analyzed by a One-way ANOVA test with original FDR method of Benjamini-Hochberg after assessment of their gaussian distribution by Shapiro-Wilk test.

**Supplementary Figure 4 related to Figure 2.**
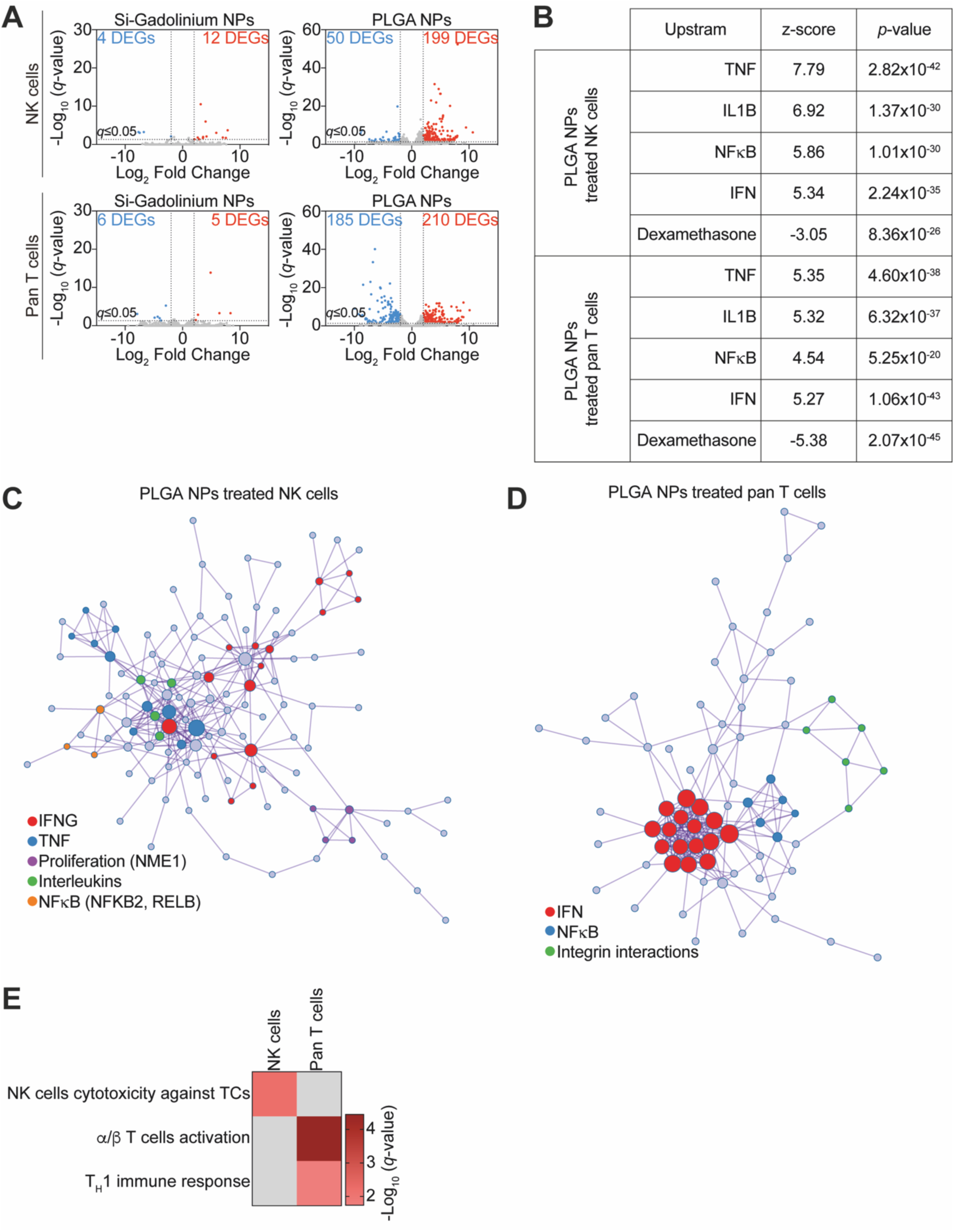
**A.** Volcano plot display differentially expressed genes between Si-Gd NPs- or PLGA NPs-treated NK cells (upper panel) or pan T cells (lower panel). Threshold -Log10 (*q*-value) ≤ 1.3 and fold-change > 2. *q*-value = adjusted *p-*value. **B.** Upstream analysis showing the top upstream molecules predicted to cause the observed gene expression changes. **C-D.** STRING predicted protein-protein interactions based on the upregulated genes in PLGA NPs-treated NK cells (**B**) and pan T cells (**C**) and their functional annotations. **E.** Heat-map showing - Log10 (*q*-value) of immune cells specific gene ontology terms assignment from upregulated genes by PLGA NPs treatment of NK cells and pan T cells.

**Supplementary Figure 5 related to Figure 2.**
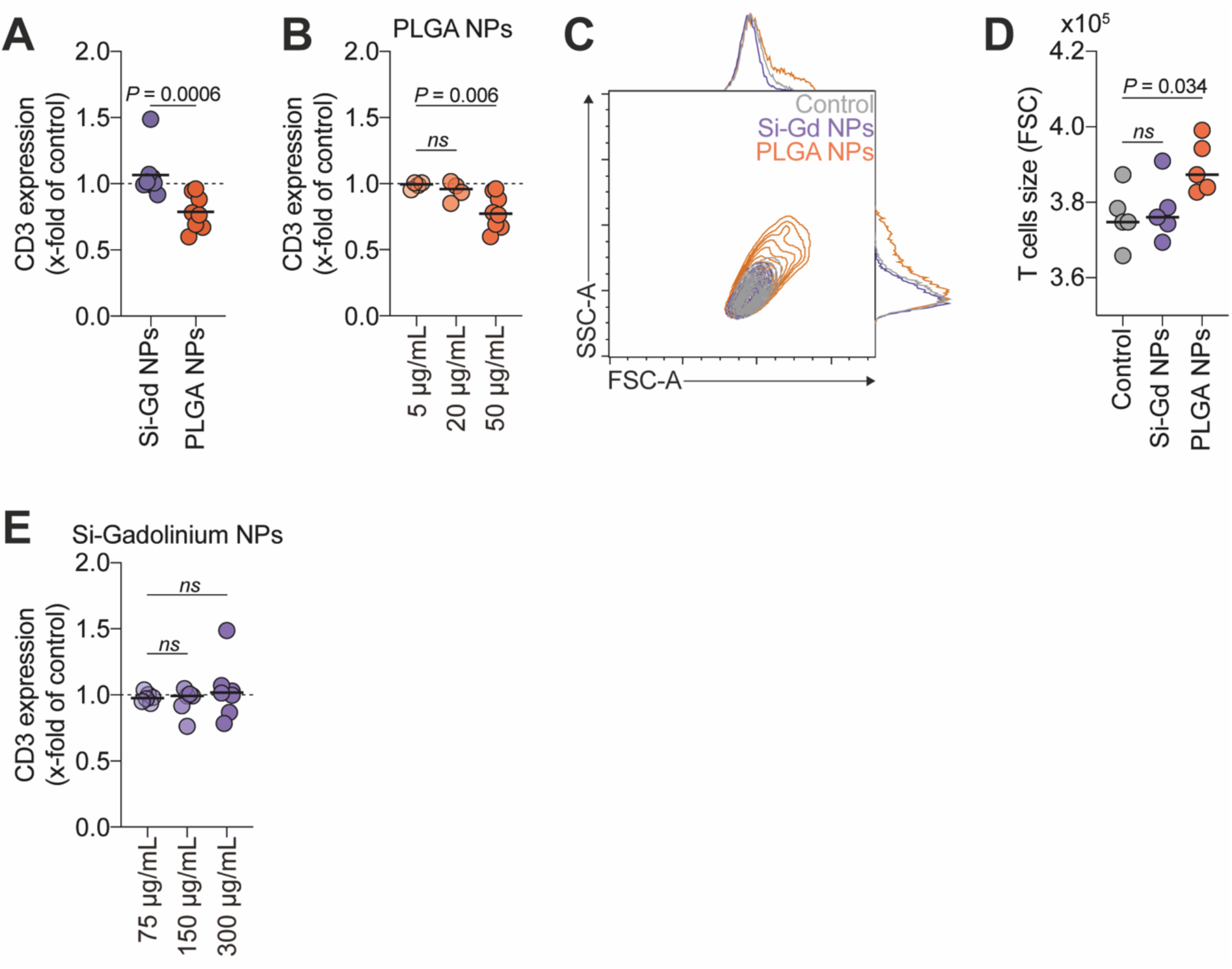
**A.** Quantification of CD3 expression at the surface of pan T cells after 48h of treatment with Si-Gd NPs or PLGA NPs. Data are represented as fold change relative to untreated control. Data are representative of 8 independent experiments and analyzed by a Mann-Whitney test after assessment of their gaussian distribution by Shapiro-Wilk test. **B.** Quantification of CD3 expression at the surface of pan T cells after 48h of treatment with increasing concentrations of PLGA NPs. Data are represented as fold change relative to untreated control. Data are representative of 4 to 8 independent experiments and analyzed by a Brown-Forsythe and Welch’s ANOVA test with original FDR method of Benjamini-Hochberg after assessment of their gaussian distribution by Shapiro-Wilk test. **C.** Representative flow cytometry contour plots of the pan T cells size (FSC) and structure (SSC) obtained by flow cytometry after Si-Gd NPs and PLGA NPs treatment of 48h. **D.** Quantification of pan T cells size based on (**C**) flow cytometry experiments after 48h of treatment with increasing concentrations of PLGA NPs. Data are representative of 5 independent experiments and analyzed by a One-way ANOVA test with original FDR method of Benjamini-Hochberg after assessment of their gaussian distribution by Shapiro-Wilk test. **D.** Quantification of CD3 expression at the surface of pan T cells after 48h of treatment with increasing concentrations of Si-Gd NPs. Data are represented as fold change relative to untreated control. Data are representative of 6 to 8 independent experiments and analyzed by a Kruskal-Wallis test after assessment of their gaussian distribution by Shapiro-Wilk test.

**Supplementary Figure 6 related to Figure 3.**
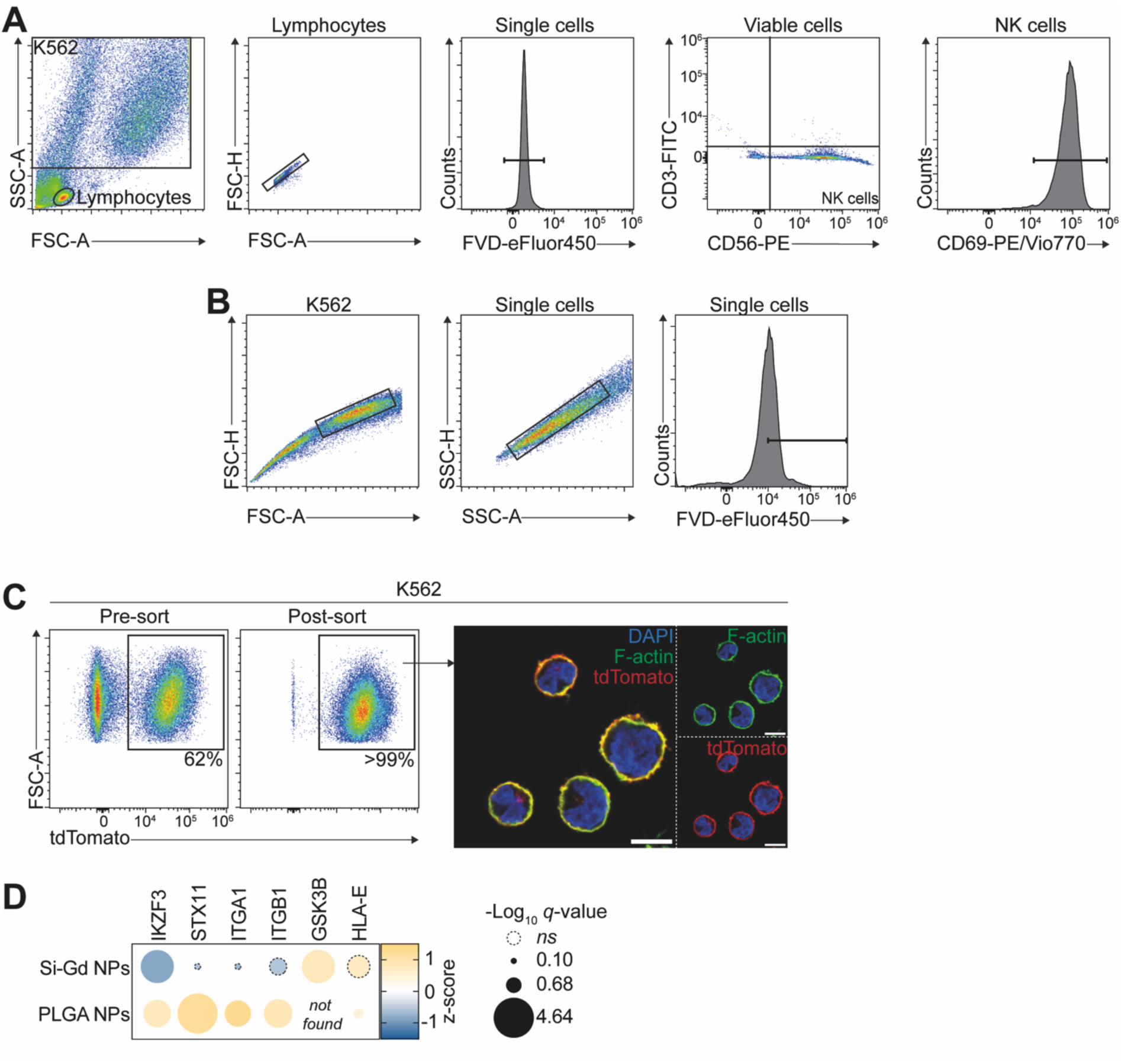
**A.** Gating strategy for assessment of NK cells activation levels. **B.** Gating strategy for assessment of NK cells-induced K562 lysis. **C.** Generation of a K562 transgenic cells expressing palmitoylated-tdTomato at their surface. Right: Representative pseudo-color flow cytometry plot before and after fluorescence-activated cells sorting for high tdTomato fluorescence. Left: Representative confocal micrographs of K562-tdTomato cells. In green Phalloidin-iFluor488, in red palmitoylated-tdTomato, in blue nuclei (DAPI). Scale bar = 10µm. **D.** Bubble plot showing Z-score values for genes/proteins expression associated with NK cells functions. Bubble size represents -Log_10_(*q*-value). Dashed lines bubbles correspond to non-significantly deregulated genes/proteins.

## Notes

### Competing Interest Statement

The authors have declared no competing interest.

## References

[1] I. Mellman, G. Coukos, G. Dranoff, Nature 2011, 480, 480.

[2] J. Galon, D. Bruni, Nat Rev Drug Discov 2019, 18, 197.

[3] L. Galluzzi, J. Humeau, A. Buqué, L. Zitvogel, G. Kroemer, Nat Rev Clin Oncol 2020, 17, 725.

[4] L. Galluzzi, M. J. Aryankalayil, C. N. Coleman, S. C. Formenti, Nat Rev Clin Oncol 2023, 20, 543.

[5] M. A. Fischbach, J. A. Bluestone, W. A. Lim, Sci. Transl. Med. 2013, 5, DOI 10.1126/scitranslmed.3005568.

[6] Z. Eshhar, T. Waks, G. Gross, D. G. Schindler, Proc. Natl. Acad. Sci. U.S.A. 1993, 90, 720.

[7] J. Maher, R. J. Brentjens, G. Gunset, I. Rivière, M. Sadelain, Nat Biotechnol 2002, 20, 70.

[8] R. J. Brentjens, J.-B. Latouche, E. Santos, F. Marti, M. C. Gong, C. Lyddane, P. D. King, S. Larson, M. Weiss, I. Rivière, M. Sadelain, Nat Med 2003, 9, 279.

[9] S. L. Maude, T. W. Laetsch, J. Buechner, S. Rives, M. Boyer, H. Bittencourt, P. Bader, M. R. Verneris, H. E. Stefanski, G. D. Myers, M. Qayed, B. De Moerloose, H. Hiramatsu, K. Schlis, K. L. Davis, P. L. Martin, E. R. Nemecek, G. A. Yanik, C. Peters, A. Baruchel, N. Boissel, F. Mechinaud, A. Balduzzi, J. Krueger, C. H. June, B. L. Levine, P. Wood, T. Taran, M. Leung, K. T. Mueller, Y. Zhang, K. Sen, D. Lebwohl, M. A. Pulsipher, S. A. Grupp, N Engl J Med 2018, 378, 439.

[10] P. Rodriguez-Otero, S. Ailawadhi, B. Arnulf, K. Patel, M. Cavo, A. K. Nooka, S. Manier, N. Callander, L. J. Costa, R. Vij, N. J. Bahlis, P. Moreau, S. R. Solomon, M. Delforge, J. Berdeja, A. Truppel-Hartmann, Z. Yang, L. Favre-Kontula, F. Wu, J. Piasecki, M. Cook, S. Giralt, N Engl J Med 2023, 388, 1002.

[11] J. N. Brudno, J. N. Kochenderfer, Blood 2016, 127, 3321.

[12] E. Liu, D. Marin, P. Banerjee, H. A. Macapinlac, P. Thompson, R. Basar, L. Nassif Kerbauy, B. Overman, P. Thall, M. Kaplan, V. Nandivada, I. Kaur, A. Nunez Cortes, K. Cao, M. Daher, C. Hosing, E. N. Cohen, P. Kebriaei, R. Mehta, S. Neelapu, Y. Nieto, M. Wang, W. Wierda, M. Keating, R. Champlin, E. J. Shpall, K. Rezvani, N Engl J Med 2020, 382, 545.

[13] T. Bald, M. F. Krummel, M. J. Smyth, K. C. Barry, Nat Immunol 2020, 21, 835.

[14] O. Demaria, L. Gauthier, G. Debroas, E. Vivier, Eur J Immunol 2021, 51, 1934.

[15] H. Salmon, K. Franciszkiewicz, D. Damotte, M.-C. Dieu-Nosjean, P. Validire, A. Trautmann, F. Mami-Chouaib, E. Donnadieu, J. Clin. Invest. 2012, 122, 899.

[16] S. Platonova, J. Cherfils-Vicini, D. Damotte, L. Crozet, V. Vieillard, P. Validire, P. André, M.-C. Dieu-Nosjean, M. Alifano, J.-F. Régnard, W.-H. Fridman, C. Sautès-Fridman, I. Cremer, Cancer Research 2011, 71, 5412.

[17] N. E. Scharping, A. V. Menk, R. S. Moreci, R. D. Whetstone, R. E. Dadey, S. C. Watkins, R. L. Ferris, G. M. Delgoffe, Immunity 2016, 45, 374.

[18] A. R. Lim, W. K. Rathmell, J. C. Rathmell, eLife 2020, 9, e55185.

[19] I. Dean, C. Y. C. Lee, Z. K. Tuong, Z. Li, C. A. Tibbitt, C. Willis, F. Gaspal, B. C. Kennedy, V. Matei-Rascu, R. Fiancette, C. Nordenvall, U. Lindforss, S. M. Baker, C. Stockmann, V. Sexl, S. A. Hammond, S. J. Dovedi, J. Mjösberg, M. R. Hepworth, G. Carlesso, M. R. Clatworthy, D. R. Withers, Nat Commun 2024, 15, 683.

[20] D. J. Slamon, B. Leyland-Jones, S. Shak, H. Fuchs, V. Paton, A. Bajamonde, T. Fleming, W. Eiermann, J. Wolter, M. Pegram, J. Baselga, L. Norton, N Engl J Med 2001, 344, 783.

[21] F. S. Hodi, S. J. O’Day, D. F. McDermott, R. W. Weber, J. A. Sosman, J. B. Haanen, R. Gonzalez, C. Robert, D. Schadendorf, J. C. Hassel, W. Akerley, A. J. M. Van Den Eertwegh, J. Lutzky, P. Lorigan, J. M. Vaubel, G. P. Linette, D. Hogg, C. H. Ottensmeier, C. Lebbé, C. Peschel, I. Quirt, J. I. Clark, J. D. Wolchok, J. S. Weber, J. Tian, M. J. Yellin, G. M. Nichol, A. Hoos, W. J. Urba, N Engl J Med 2010, 363, 711.

[22] F. Berner, D. Bomze, S. Diem, O. H. Ali, M. Fässler, S. Ring, R. Niederer, C. J. Ackermann, P. Baumgaertner, N. Pikor, C. G. Cruz, W. Van De Veen, M. Akdis, S. Nikolaev, H. Läubli, A. Zippelius, F. Hartmann, H.-W. Cheng, G. Hönger, M. Recher, J. Goldman, A. Cozzio, M. Früh, J. Neefjes, C. Driessen, B. Ludewig, A. N. Hegazy, W. Jochum, D. E. Speiser, L. Flatz, JAMA Oncol 2019, 5, 1043.

[23] D. B. Johnson, J. M. Balko, M. L. Compton, S. Chalkias, J. Gorham, Y. Xu, M. Hicks, I. Puzanov, M. R. Alexander, T. L. Bloomer, J. R. Becker, D. A. Slosky, E. J. Phillips, M. A. Pilkinton, L. Craig-Owens, N. Kola, G. Plautz, D. S. Reshef, J. S. Deutsch, R. P. Deering, B. A. Olenchock, A. H. Lichtman, D. M. Roden, C. E. Seidman, I. J. Koralnik, J. G. Seidman, R. D. Hoffman, J. M. Taube, L. A. Diaz, R. A. Anders, J. A. Sosman, J. J. Moslehi, N Engl J Med 2016, 375, 1749.

[24] V. Mittelheisser, M. Banerjee, X. Pivot, L. J. Charbonnière, J. Goetz, A. Detappe, Adv. Therap. 2020, 3, 2000134.

[25] T. R. Fadel, M. Look, P. A. Staffier, G. L. Haller, L. D. Pfefferle, T. M. Fahmy, Langmuir 2010, 26, 5645.

[26] T. R. Fadel, F. A. Sharp, N. Vudattu, R. Ragheb, J. Garyu, D. Kim, E. Hong, N. Li, G. L. Haller, L. D. Pfefferle, S. Justesen, K. C. Herold, T. M. Fahmy, Nature Nanotech 2014, 9, 639.

[27] D. Schmid, C. G. Park, C. A. Hartl, N. Subedi, A. N. Cartwright, R. B. Puerto, Y. Zheng, J. Maiarana, G. J. Freeman, K. W. Wucherpfennig, D. J. Irvine, M. S. Goldberg, Nat Commun 2017, 8, 1747.

[28] T. T. Smith, S. B. Stephan, H. F. Moffett, L. E. McKnight, W. Ji, D. Reiman, E. Bonagofski, M. E. Wohlfahrt, S. P. S. Pillai, M. T. Stephan, Nature Nanotech 2017, 12, 813.

[29] N. N. Parayath, S. B. Stephan, A. L. Koehne, P. S. Nelson, M. T. Stephan, Nat Commun 2020, 11, 6080.

[30] L. Carmès, G. Bort, F. Lux, L. Seban, P. Rocchi, Z. Muradova, A. Hagège, L. Heinrich-Balard, F. Delolme, V. Gueguen-Chaignon, C. Truillet, S. Crowley, E. Bello, T. Doussineau, M. Dougan, O. Tillement, J. D. Schoenfeld, N. Brown, R. Berbeco, Nanoscale 2024, 16, 2347.

[31] A. Mignot, C. Truillet, F. Lux, L. Sancey, C. Louis, F. Denat, F. Boschetti, L. Bocher, A. Gloter, O. Stéphan, R. Antoine, P. Dugourd, D. Luneau, G. Novitchi, L. C. Figueiredo, P. C. de Morais, L. Bonneviot, B. Albela, F. Ribot, L. Van Lokeren, I. Déchamps-Olivier, F. Chuburu, G. Lemercier, C. Villiers, P. N. Marche, G. Le Duc, S. Roux, O. Tillement, P. Perriat, Chemistry A European J 2013, 19, 6122.

[32] V.-L. Tran, V. Thakare, F. Rossetti, A. Baudouin, G. Ramniceanu, B.-T. Doan, N. Mignet, C. Comby-Zerbino, R. Antoine, P. Dugourd, F. Boschetti, F. Denat, C. Louis, S. Roux, T. Doussineau, O. Tillement, F. Lux, J. Mater. Chem. B 2018, 6, 4821.

[33] T. Lammers, Advanced Materials 2024, 36, 2312169.

[34] B. Vis, R. E. Hewitt, T. P. Monie, C. Fairbairn, S. D. Turner, S. D. Kinrade, J. J. Powell, Proc. Natl. Acad. Sci. U.S.A. 2020, 117, 285.

[35] S. Weller, X. Li, L. R. Petersen, P. Kempen, G. Clergeaud, T. L. Andresen, International Immunopharmacology 2024, 129, 111643.

[36] M. Vitale, C. Bottino, S. Sivori, L. Sanseverino, R. Castriconi, E. Marcenaro, R. Augugliaro, L. Moretta, A. Moretta, The Journal of Experimental Medicine 1998, 187, 2065.

[37] O. Dufva, S. Gandolfi, J. Huuhtanen, O. Dashevsky, H. Duàn, K. Saeed, J. Klievink, P. Nygren, J. Bouhlal, J. Lahtela, A. Näätänen, B. R. Ghimire, T. Hannunen, P. Ellonen, H. Lähteenmäki, P. Rumm, J. Theodoropoulos, E. Laajala, J. Härkönen, P. Pölönen, M. Heinäniemi, M. Hollmén, S. Yamano, R. Shirasaki, D. A. Barbie, J. A. Roth, R. Romee, M. Sheffer, H. Lähdesmäki, D. A. Lee, R. De Matos Simoes, M. Kankainen, C. S. Mitsiades, S. Mustjoki, Immunity 2023, 56, 2816.

[38] A. Pfefferle, B. Jacobs, H. Netskar, E. H. Ask, S. Lorenz, T. Clancy, J. P. Goodridge, E. Sohlberg, K.-J. Malmberg, Cell Reports 2019, 29, 2284.

[39] D. Urlaub, K. Höfer, M.-L. Müller, C. Watzl, The Journal of Immunology 2017, 198, 1944.

[40] J. P. Böttcher, E. Bonavita, P. Chakravarty, H. Blees, M. Cabeza-Cabrerizo, S. Sammicheli, N. C. Rogers, E. Sahai, S. Zelenay, C. Reis E Sousa, Cell 2018, 172, 1022.

[41] T. D. Holmes, R. V. Pandey, E. Y. Helm, H. Schlums, H. Han, T. M. Campbell, T. T. Drashansky, S. Chiang, C.-Y. Wu, C. Tao, M. Shoukier, E. Tolosa, S. Von Hardenberg, M. Sun, C. Klemann, R. A. Marsh, C. M. Lau, Y. Lin, J. C. Sun, R. Månsson, F. Cichocki, D. Avram, Y. T. Bryceson, Sci. Immunol. 2021, 6, eabc9801.

[42] E. K. Santosa, H. Kim, T. Rückert, J.-B. Le Luduec, A. J. Abbasi, C. K. Wingert, L. Peters, J. N. Frost, K. C. Hsu, C. Romagnani, J. C. Sun, Nat Immunol 2023, 24, 1685.

[43] G. Min-Oo, N. A. Bezman, S. Madera, J. C. Sun, L. L. Lanier, Journal of Experimental Medicine 2014, 211, 1289.

[44] R. B. Mailliard, S. M. Alber, H. Shen, S. C. Watkins, J. M. Kirkwood, R. B. Herberman, P. Kalinski, The Journal of Experimental Medicine 2005, 202, 941.

[45] P. Lee, T. Yamada, C. S. Park, Y. Shen, M. Puppi, H. D. Lacorazza, Immunol Cell Biol 2015, 93, 605.

[46] P. A. Szabo, H. M. Levitin, M. Miron, M. E. Snyder, T. Senda, J. Yuan, Y. L. Cheng, E. C. Bush, P. Dogra, P. Thapa, D. L. Farber, P. A. Sims, Nat Commun 2019, 10, 4706.

[47] L. Loyal, S. Warth, K. Jürchott, F. Mölder, C. Nikolaou, N. Babel, M. Nienen, S. Durlanik, R. Stark, B. Kruse, M. Frentsch, R. Sabat, K. Wolk, A. Thiel, Nat Commun 2020, 11, 6357.

[48] X.-Y. Li, D. Corvino, B. Nowlan, A. R. Aguilera, S. S. Ng, M. Braun, A. R. Cillo, T. Bald, M. J. Smyth, C. R. Engwerda, Cancer Immunology Research 2022, 10, 154.

[49] M. Kurachi, R. A. Barnitz, N. Yosef, P. M. Odorizzi, M. A. DiIorio, M. E. Lemieux, K. Yates, J. Godec, M. G. Klatt, A. Regev, E. J. Wherry, W. N. Haining, Nat Immunol 2014, 15, 373.

[50] Z. Qiu, C. Khairallah, G. Romanov, B. S. Sheridan, The Journal of Immunology 2020, 205, 901.

[51] K. J. Oestreich, A. S. Weinmann, Current Opinion in Immunology 2012, 24, 191.

[52] Y. Liu, R. Bockermann, M. Hadi, I. Safari, B. Carrion, M. Kveiborg, S. Issazadeh-Navikas, Cell Mol Immunol 2021, 18, 1904.

[53] U. Gubler, A. O. Chua, D. S. Schoenhaut, C. M. Dwyer, W. McComas, R. Motyka, N. Nabavi, A. G. Wolitzky, P. M. Quinn, P. C. Familletti, Proc. Natl. Acad. Sci. U.S.A. 1991, 88, 4143.

[54] G. S. Kelner, J. Kennedy, K. B. Bacon, S. Kleyensteuber, D. A. Largaespada, N. A. Jenkins, N. G. Copeland, J. F. Bazan, K. W. Moore, T. J. Schall, A. Zlotnik, Science 1994, 266, 1395.

[55] A. J. Ozga, M. T. Chow, A. D. Luster, Immunity 2021, 54, 859.

[56] R. Lachmann, M. Bajwa, S. Vita, H. Smith, E. Cheek, A. Akbar, F. Kern, J Virol 2012, 86, 1001.

[57] W. M. Schneider, M. D. Chevillotte, C. M. Rice, Annu. Rev. Immunol. 2014, 32, 513.

[58] E. Cano-Gamez, B. Soskic, T. I. Roumeliotis, E. So, D. J. Smyth, M. Baldrighi, D. Willé, N. Nakic, J. Esparza-Gordillo, C. G. C. Larminie, P. G. Bronson, D. F. Tough, W. C. Rowan, J. S. Choudhary, G. Trynka, Nat Commun 2020, 11, 1801.

[59] A. L. Symonds, W. Zheng, T. Miao, H. Wang, T. Wang, R. Kiome, X. Hou, S. Li, P. Wang, Life Sci. Alliance 2020, 3, e202000766.

[60] A. L. J. Symonds, T. Miao, Z. Busharat, S. Li, P. Wang, Cancer Immunol Immunother 2023, 72, 1139.

[61] M. Binnewies, E. W. Roberts, K. Kersten, V. Chan, D. F. Fearon, M. Merad, L. M. Coussens, D. I. Gabrilovich, S. Ostrand-Rosenberg, C. C. Hedrick, R. H. Vonderheide, M. J. Pittet, R. K. Jain, W. Zou, T. K. Howcroft, E. C. Woodhouse, R. A. Weinberg, M. F. Krummel, Nat Med 2018, 24, 541.

[62] M. L. Holmes, N. D. Huntington, R. P. Thong, J. Brady, Y. Hayakawa, C. E. Andoniou, P. Fleming, W. Shi, G. K. Smyth, M. A. Degli-Esposti, G. T. Belz, A. Kallies, S. Carotta, M. J. Smyth, S. L. Nutt, The EMBO Journal 2014, 33, 2721.

[63] E. Hegewisch-Solloa, S. Seo, B. L. Mundy-Bosse, A. Mishra, E. H. Waldman, S. Maurrasse, E. Grunstein, T. J. Connors, A. G. Freud, E. M. Mace, The Journal of Immunology 2021, 207, 950.

[64] O. D’Orlando, F. Zhao, B. Kasper, Z. Orinska, J. Müller, I. Hermans-Borgmeyer, G. M. Griffiths, U. Zur Stadt, S. Bulfone-Paus, Eur J Immunol 2013, 43, 194.

[65] E. Le Dréan, F. Vély, L. Olcese, A. Cambiaggi, S. Guia, G. Krystal, N. Gervois, A. Moretta, F. Jotereau, E. Vivier, Eur. J. Immunol. 1998, 28, 264.

[66] R. Parameswaran, P. Ramakrishnan, S. A. Moreton, Z. Xia, Y. Hou, D. A. Lee, K. Gupta, M. deLima, R. C. Beck, D. N. Wald, Nat Commun 2016, 7, 11154.

[67] F. Cichocki, B. Valamehr, R. Bjordahl, B. Zhang, B. Rezner, P. Rogers, S. Gaidarova, S. Moreno, K. Tuininga, P. Dougherty, V. McCullar, P. Howard, D. Sarhan, E. Taras, H. Schlums, S. Abbot, D. Shoemaker, Y. T. Bryceson, B. R. Blazar, S. Wolchko, S. Cooley, J. S. Miller, Cancer Research 2017, 77, 5664.

[68] Y. Kashii, R. Giorda, R. B. Herberman, T. L. Whiteside, N. L. Vujanovic, J Immunol 1999, 163, 5358.

[69] G. Z. Tau, S. N. Cowan, J. Weisburg, N. S. Braunstein, P. B. Rothman, The Journal of Immunology 2001, 167, 5574.

[70] V. P. Patel, M. Moran, T. A. Low, M. C. Miceli, The Journal of Immunology 2001, 166, 754.

[71] D. M. Desai, M. E. Newton, T. Kadlecek, A. Weiss, Nature 1990, 348, 66.

[72] J. L. Pomerantz, E. M. Denny, D. Baltimore, EMBO J 2002, 21, 5184.

[73] K. Chemin, A. Bohineust, S. Dogniaux, M. Tourret, S. Guégan, F. Miro, C. Hivroz, The Journal of Immunology 2012, 189, 2159.

[74] A. S. Zhovmer, A. Manning, C. Smith, A. Nguyen, O. Prince, P. J. Sáez, X. Ma, D. Tsygankov, A. X. Cartagena-Rivera, N. A. Singh, R. K. Singh, E. D. Tabdanov, Sci. Adv. 2024, 10, eadi1788.

[75] A. Roumier, J. C. Olivo-Marin, M. Arpin, F. Michel, M. Martin, P. Mangeat, O. Acuto, A. Dautry-Varsat, A. Alcover, Immunity 2001, 15, 715.

[76] J. C. Nolz, T. S. Gomez, P. Zhu, S. Li, R. B. Medeiros, Y. Shimizu, J. K. Burkhardt, B. D. Freedman, D. D. Billadeau, Current Biology 2006, 16, 24.

[77] T. S. Gomez, K. Kumar, R. B. Medeiros, Y. Shimizu, P. J. Leibson, D. D. Billadeau, Immunity 2007, 26, 177.

[78] B. P. Nicolet, A. Guislain, F. P. J. Van Alphen, R. Gomez-Eerland, T. N. M. Schumacher, M. Van Den Biggelaar, M. C. Wolkers, Proc. Natl. Acad. Sci. U.S.A. 2020, 117, 6686.

[79] B. P. Nicolet, A. Guislain, M. C. Wolkers, The Journal of Immunology 2021, 207, 2966.

[80] Z. Wang, J. Shang, Y. Qiu, H. Cheng, M. Tao, E. Xie, X. Pei, W. Li, L. Zhang, A. Wu, G. Li, Cell Reports 2024, 43, 113796.

[81] M. Kurowska, N. Goudin, N. T. Nehme, M. Court, J. Garin, A. Fischer, G. De Saint Basile, G. Ménasché, Blood 2012, 119, 3879.

[82] V. Das, B. Nal, A. Dujeancourt, M.-I. Thoulouze, T. Galli, P. Roux, A. Dautry-Varsat, A. Alcover, Immunity 2004, 20, 577.

[83] M. R. Marshall, V. Pattu, M. Halimani, M. Maier-Peuschel, M.-L. Müller, U. Becherer, W. Hong, M. Hoth, T. Tschernig, Y. T. Bryceson, J. Rettig, Journal of Cell Biology 2015, 210, 135.

[84] M. L. Dustin, Cancer Immunology Research 2014, 2, 1023.

[85] I. Gramaglia, A. D. Weinberg, M. Lemon, M. Croft, J Immunol 1998, 161, 6510.

[86] G. Verdeil, D. Puthier, C. Nguyen, A.-M. Schmitt-Verhulst, N. Auphan-Anezin, The Journal of Immunology 2006, 176, 4834.

[87] M. Pescatori, D. Bedognetti, E. Venturelli, C. Ménard-Moyon, C. Bernardini, E. Muresu, A. Piana, G. Maida, R. Manetti, F. Sgarrella, A. Bianco, L. G. Delogu, Biomaterials 2013, 34, 4395.

[88] N. Kumbhojkar, S. Prakash, T. Fukuta, K. Adu-Berchie, N. Kapate, R. An, S. Darko, V. Chandran Suja, K. S. Park, A. P. Gottlieb, M. G. Bibbey, M. Mukherji, L. L.-W. Wang, D. J. Mooney, S. Mitragotri, Nat. Biomed. Eng 2024, 8, 579.

[89] T. Walzer, M. Dalod, S. H. Robbins, L. Zitvogel, E. Vivier, Blood 2005, 106, 2252.

[90] C. Verry, S. Dufort, B. Lemasson, S. Grand, J. Pietras, I. Troprès, Y. Crémillieux, F. Lux, S. Mériaux, B. Larrat, J. Balosso, G. Le Duc, E. L. Barbier, O. Tillement, Sci. Adv. 2020, 6, eaay5279.

[91] C. Verry, S. Dufort, J. Villa, M. Gavard, C. Iriart, S. Grand, J. Charles, B. Chovelon, J.-L. Cracowski, J.-L. Quesada, C. Mendoza, L. Sancey, A. Lehmann, F. Jover, J.-Y. Giraud, F. Lux, Y. Crémillieux, S. McMahon, P. J. Pauwels, D. Cagney, R. Berbeco, A. Aizer, E. Deutsch, M. Loeffler, G. Le Duc, O. Tillement, J. Balosso, Radiotherapy and Oncology 2021, 160, 159.

[92] M. E. Rodriguez-Ruiz, I. Vitale, K. J. Harrington, I. Melero, L. Galluzzi, Nat Immunol 2020, 21, 120.

[93] B. Diskin, S. Adam, M. F. Cassini, G. Sanchez, M. Liria, B. Aykut, C. Buttar, E. Li, B. Sundberg, R. D. Salas, R. Chen, J. Wang, M. Kim, M. S. Farooq, S. Nguy, C. Fedele, K. H. Tang, T. Chen, W. Wang, M. Hundeyin, J. A. K. Rossi, E. Kurz, M. I. U. Haq, J. Karlen, E. Kruger, Z. Sekendiz, D. Wu, S. A. A. Shadaloey, G. Baptiste, G. Werba, S. Selvaraj, C. Loomis, K.-K. Wong, J. Leinwand, G. Miller, Nat Immunol 2020, 21, 442.

[94] W. Piao, L. Li, V. Saxena, J. Iyyathurai, R. Lakhan, Y. Zhang, I. T. Lape, C. Paluskievicz, K. L. Hippen, Y. Lee, E. Silverman, M. W. Shirkey, L. V. Riella, B. R. Blazar, J. S. Bromberg, Nat Commun 2022, 13, 2176.

[95] L. García-Hevia, R. Soltani, J. González, O. Chaloin, C. Ménard-Moyon, A. Bianco, M. L. Fanarraga, Bioactive Materials 2024, 34, 237.

[96] C. Charpentier, V. Cifliku, J. Goetz, A. Nonat, C. Cheignon, M. Cardoso Dos Santos, L. Francés-Soriano, K. Wong, L. J. Charbonnière, N. Hildebrandt, Chemistry A European J 2020, 26, 14602.

[97] L. Zhang, J. M. Chan, F. X. Gu, J.-W. Rhee, A. Z. Wang, A. F. Radovic-Moreno, F. Alexis, R. Langer, O. C. Farokhzad, ACS Nano 2008, 2, 1696.

[98] S. Andrews, “FastQC: a quality control tool for high throughput sequence data.,” can be found under https://www.bioinformatics.babraham.ac.uk/projects/fastqc/, 2010.

[99] A. Dobin, C. A. Davis, F. Schlesinger, J. Drenkow, C. Zaleski, S. Jha, P. Batut, M. Chaisson, T. R. Gingeras, Bioinformatics 2013, 29, 15.

[100] S. Anders, P. T. Pyl, W. Huber, Bioinformatics 2015, 31, 166.

[101] M. I. Love, W. Huber, S. Anders, Genome Biol 2014, 15, 550.

[102] R. C. Gentleman, V. J. Carey, D. M. Bates, B. Bolstad, M. Dettling, S. Dudoit, B. Ellis, L. Gautier, Y. Ge, J. Gentry, K. Hornik, T. Hothorn, W. Huber, S. Iacus, R. Irizarry, F. Leisch, C. Li, M. Maechler, A. J. Rossini, G. Sawitzki, C. Smith, G. Smyth, L. Tierney, J. Y. Yang, J. Zhang, Genome Biol 2004, 5, R80.

[103] Y. Zhou, B. Zhou, L. Pache, M. Chang, A. H. Khodabakhshi, O. Tanaseichuk, C. Benner, S. K. Chanda, Nat Commun 2019, 10, 1523.

[104] C. S. Hughes, S. Moggridge, T. Müller, P. H. Sorensen, G. B. Morin, J. Krijgsveld, Nat Protoc 2019, 14, 68.

[105] D. Bouyssié, A.-M. Hesse, E. Mouton-Barbosa, M. Rompais, C. Macron, C. Carapito, A. Gonzalez De Peredo, Y. Couté, V. Dupierris, A. Burel, J.-P. Menetrey, A. Kalaitzakis, J. Poisat, A. Romdhani, O. Burlet-Schiltz, S. Cianférani, J. Garin, C. Bruley, Bioinformatics 2020, 36, 3148.

[106] S. Wieczorek, F. Combes, C. Lazar, Q. Giai Gianetto, L. Gatto, A. Dorffer, A.-M. Hesse, Y. Couté, M. Ferro, C. Bruley, T. Burger, Bioinformatics 2017, 33, 135.

[107] V. P. Sharma, B. Tang, Y. Wang, C. L. Duran, G. S. Karagiannis, E. A. Xue, D. Entenberg, L. Borriello, A. Coste, R. J. Eddy, G. Kim, X. Ye, J. G. Jones, E. Grunblatt, N. Agi, S. Roy, G. Bandyopadhyaya, E. Adler, C. R. Surve, D. Esposito, S. Goswami, J. E. Segall, W. Guo, J. S. Condeelis, L. M. Wakefield, M. H. Oktay, Nat Commun 2021, 12, 7300.

[108] Y. Perez-Riverol, J. Bai, C. Bandla, D. García-Seisdedos, S. Hewapathirana, S. Kamatchinathan, D. J. Kundu, A. Prakash, A. Frericks-Zipper, M. Eisenacher, M. Walzer, S. Wang, A. Brazma, J. A. Vizcaíno, Nucleic Acids Research 2022, 50, D543.

